# A general platform for targeting MHC-II antigens via a single loop

**DOI:** 10.1101/2024.01.26.577489

**Authors:** Haotian Du, Jingjia Liu, Kevin M. Jude, Xinbo Yang, Ying Li, Braxton Bell, Hongli Yang, Audrey Kassardjian, Ali Mobedi, Udit Parekh, R. Andres Parra Sperberg, Jean-Philippe Julien, Elizabeth D. Mellins, K. Christopher Garcia, Po-Ssu Huang

## Abstract

Class-II major histocompatibility complexes (MHC-IIs) are central to the communications between CD4+ T cells and antigen presenting cells (APCs), but intrinsic structural features associated with MHC-II make it difficult to develop a general targeting system with high affinity and antigen specificity. Here, we introduce a protein platform, Targeted Recognition of Antigen-MHC Complex Reporter for MHC-II (*TRACeR-II*), to enable the rapid development of peptide-specific MHC-II binders. *TRACeR-II* has a small helical bundle scaffold and uses an unconventional mechanism to recognize antigens via a single loop. This unique antigen-recognition mechanism renders this platform highly versatile and amenable to direct structural modeling of the interactions with the antigen. We demonstrate that *TRACeR-II* binders can be rapidly evolved across multiple alleles, while computational protein design can produce specific binding sequences for a SARS-CoV-2 peptide of unknown complex structure. *TRACeR-II* sheds light on a simple and straightforward approach to address the MHC peptide targeting challenge, without relying on combinatorial selection on complementarity determining region (CDR) loops. It presents a promising basis for further exploration in immune response modulation as well as a broad range of theragnostic applications.

## Introduction

T cell immunity relies on interactions between major histocompatibility complexes (MHCs) and T cell receptors (TCRs) to trigger an antigen-dependent response, with presentation of antigen on MHCs a central aspect of the process. Class-II MHCs, which are found on antigen presenting cells (APCs) and other cell types including transformed cells^1^, play a significant role in the detection of foreign antigens^2^, in driving autoimmune diseases^3, 4^, and potentially in mediating microbiome tolerance^5, 6^. Although recent studies have shed light on the relationship between APCs, MHC-II and antigen presentation^7, 8^, much of the fundamental biology and theragnostic potential remains to be explored.

Presently, TCRs and TCR-like antibodies are the only approaches for targeting peptide-MHC complexes (pMHCs)^9-13^. However, the development of TCRs or antibodies require *de novo* screening for each pMHC target with no guaranteed success. Due to the flat antigen structure on MHC-IIs, there is few structural features for binders to engage. To achieve discrimination of diverse antigens, the canonical TCR docking pose on MHC-II allows as many as 4 CDRs to align along the antigen^14,15^. The use of flexible loops potentially causes the lower intrinsic affinity between TCR and pMHC-II than pMHC-I^16^, making initial binder identification even more difficult. Additionally, loop-based engineering of TCRs and TCR-like antibodies from directed evolution can adopt diverse binding modes, making it hard to select for a specific binding pose during subsequent development process. Because of these issues, the engineering of TCRs and TCR-like antibodies often requires large combinatorial libraries and extensive screening efforts^17-19^.

Having a molecular platform for facile and rapid development of antigen-focused MHC-II binders will have a significant impact in a variety of biotechnological and biomedical applications. While recent advances in protein structure prediction and design have advanced our understanding and engineering capabilities of protein-protein interactions^20, 21^, they have not contributed significantly to MHC-II binder engineering due to the intricate structural complexity of TCR, antigen and MHC-II.

All MHC-II molecules share a common backbone topology, and MHC-II antigens reside in a highly confined structural space. This inspired us to explore alternative mechanisms to employ non-TCR (nor antibody) structures. We hypothesize that the complexity of the MHC-II targeting problem can be significantly reduced by defining a constant binding mode and restricting the region in the binder structure malleable to the antigen to one segment. We reason that as a general MHC targeting platform, the proposed design should consider two essential elements: first, an MHC binding element to bind specifically to MHC and not other surface receptors and ideally should be compatible with different MHC alleles to address polymorphism (Fig. 1a); second, a peptide recognition element to complement the peptide of interest, conveying target specificity. Inspired by the unique binding mode of a superantigen, *Mycoplasma arthritidis* mitogen (MAM)^22^, we created a protein platform that exquisitely satisfies these two requirements.

**Figure 1.**
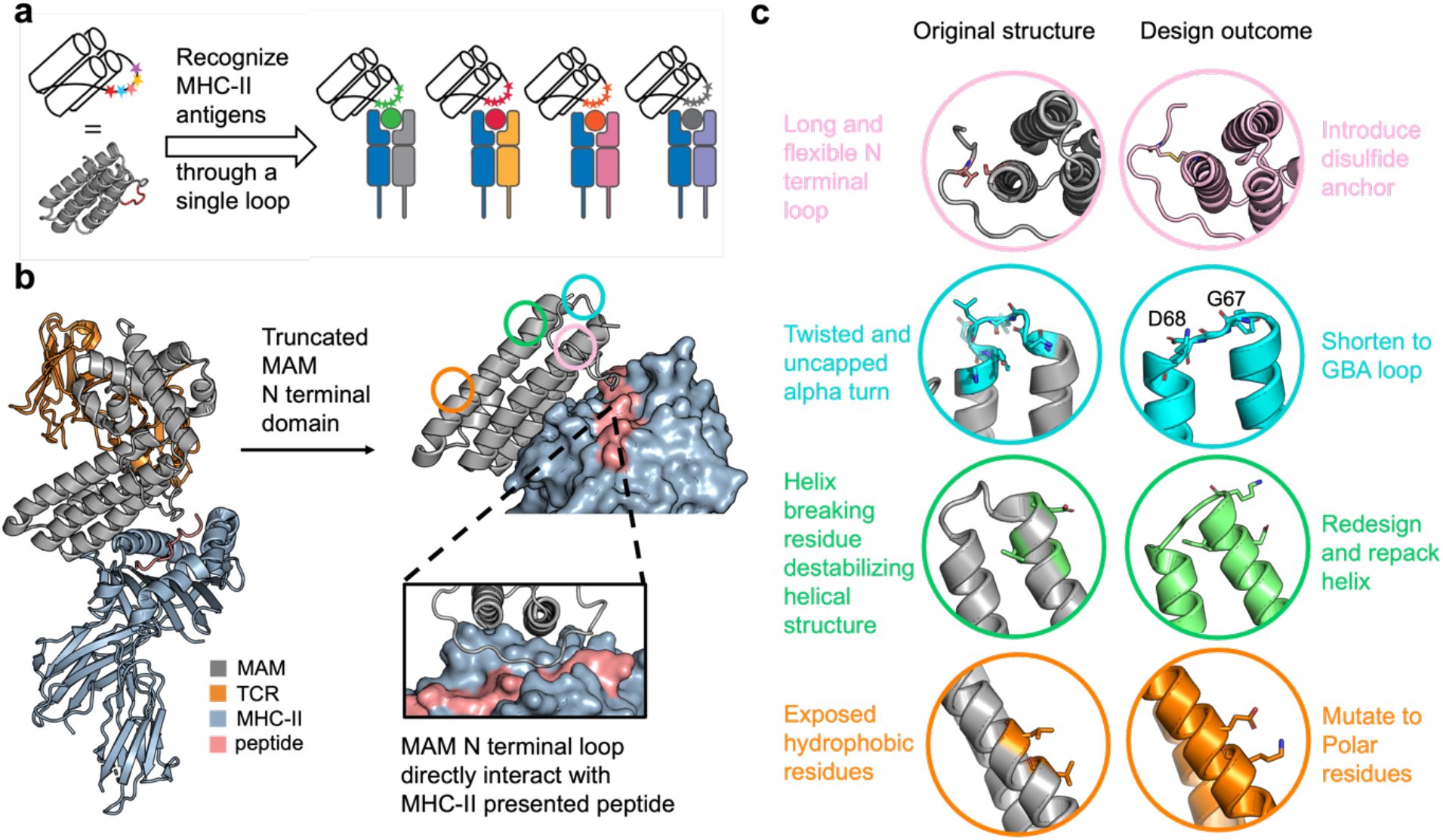
Design scheme of *TRACeR-II* platform. **a**. Facile development of peptide-focused MHC-II binders by mutating a single adaptive loop. **b**. Left: Ternary complex structure of the MAM-MHC-TCR complex (PDB ID: 2ICW). Right: The binding structure between truncated MAM N terminal domain and pMHC-II. **c**. Structural optimization by RosettaRemodel.

This single-chained, stable, and versatile platform is named Targeted Recognition of Antigen-MHC Complex Reporter for MHC-II, or *TRACeR-II. TRACeR-II* can be applied across a range of MHC-II alleles. Most importantly, *TRACeR-II’s* molecular recognition mechanism utilizes a single adaptable loop to achieve antigen specificity, and this enables rational *in silico* design of *TRACeRs* against specific antigen-of-interest directly. This unprecedented capability has the potential to serve as a solution to leverage the vast expanding immunopeptidomic data.

## Results

### Molecular engineering inspired by superantigen MAM

Superantigens are typically produced by bacteria, viruses, and other microorganisms. They broadly activate T cells by bridging MHC-II and TCR, overwriting antigen restriction between an antigen presenting cell and a T cell^23^. MAM is a superantigen with a unique binding mode to the MHC-II (Protein Data Bank (PDB) ID: 2ICW^22^) featuring an L-shaped structure consisting of two domains. Its N-terminal domain contacts pMHC-II, while both N- and C-terminal domains jointly form the interface with TCR. The native structure also has a flexible loop (residues 1-20) which is located above the antigen binding groove on MHC-II. (Fig. 1b)

The flexible loop is conveniently located above the antigen in the MHC-II groove, but despite the loop’s proximity, native MAM is agnostic to the antigen and can bind to different MHC subtypes^24^. MAM’s unique structure led us to consider the possibility of engineering MAM as a general binding platform for targeting pMHC-II in a peptide-specific manner (Fig. 1a). Additionally, MAM’s compatibility with multiple HLA alleles suggests a possible mechanism for multi-allelic, antigen-focused binding^24^.

Deleting the flexible loop from MAM completely abolishes its pMHC binding (Supplementary Fig. S1), revealing truly surprising aspects of the MAM structure. In the unbound MAM crystal structure (PDB ID: 3KPH), the first 15 residues are too labile to be resolved, so their critical contribution to binding is unusual. We had expected residual binding signals from the helical bundle alone, but the complete loss of binding makes MAM an even more attractive scaffold. It appears that the loop is the critical element that allows MAM to make a crowbar-like shape to saddle over MHC-II’s α1 helix, with the loop being the prying tip. That MAM has no significant affinity to MHC-II without the loop would make any positive binding signal truly antigen-focused.

Encouraged by these observations, we undertook extensive engineering efforts to transform MAM into an MHC-antigen binding platform. When developing a superantigen into a targeting platform, the intrinsic toxicity from engaging T cells is of primary concern. MAM can be rendered inert by eliminating its C-terminal domain, which abolishes MAM’s ability to bind TCR, and the N-terminal domain alone is sufficient to bind pMHC-II (Supplementary Fig. S2).

In our design scheme, we planned to turn the N-terminal flexible loop into the antigen recognition element (ARE). To develop it into a segment that can accommodate a wide range of possible antigens, the natively disordered loop would need to be anchored at the free end to tolerate the introduction of randomized mutations in this region. To achieve this, we computationally designed a specific disulfide bond that anchors the loop to the helical bundle scaffold (Fig. 1c, line 1). This modification involving residues 5 and 119 simultaneously reduced loop flexibility and improved overall stability. The design was incorporated into the *TRACeR-II* scaffold (Supplementary Fig. S3).

To further optimize the structure, we identified a few non-ideal structural features associated with the initial helical bundle and computationally redesigned these regions with RosettaRemodel^25^ (Fig. 1c, line 2-4). First, we redesigned one of the alpha-alpha turns. The native alpha-alpha turn between the second and third helix has a non-ideal loop connection, with a twisted geometry and a missing N-cap of the third helix. We redesigned this turn into a more energetically favorable GBA loop^26^, featuring a Gly67 break in the second helix and a Asp68 cap on the third helix (Fig 1c, line 2). Our experimental results confirmed that this redesign resulted in more than 100-fold increase in recombinant protein production yield, denoting a huge improvement on scaffold stability. We also addressed several other stability issues, including the mutation of G61D with repacking of surrounding residues to reduce the glycine’s helix-breaking effect (Fig. 1c, line 3), and the L47K and L50E mutations to reduce solvent-exposed hydrophobicity (Fig. 1d, line 4). The immunogenicity of the optimized helical bundle was evaluated in mice through multiple injections over a two-week period. No signs of toxicity or immunogenicity were observed based on no changes in body weight and no notable development of antibody titers, respectively, during this study duration (Supplementary Fig. S4).

### Discrimination of peptides presented by HLA-DR1

Once an optimal *TRACeR* starting scaffold had been established, we applied a combinatorial library approach to adapt the ARE region to recognize diverse peptides. We selected 5 continuous positions on the ARE region that are closest to MHC-II-bound peptide based on the crystal structure and built a yeast library using degenerate codons (NNK) with a final diversity of 3.2 × 10^6^ (Supplementary Table 1). We built *TRACeRs* for three different MHC-II antigens on HLA-DR1 to demonstrate the feasibility of this binding platform. These targets encompass viral (HLA-DR1(DRA1*01:01/DRB1*01:01)/HA307-319 PKYVKQNTLKLAT), cancer (HLA-DR1/NYESO187-101 LLEFYLAMPFATPME^27^), and self (HLA-DR1/CLIP87-101 PVSKMRMATPLLMQA) derived antigens. The yeast library was screened with fluorescently labeled MHC-II tetramers. Affinity and specificity selection were conducted simultaneously with fluorescence-activated cell sorting (FACS). We isolated *TRACeRs* from the same pool that specifically bound cognate targets without cross-reacting with the other two (Supplementary Fig. S5). Loop sequences were converged to completely distinct sequence patterns for each of the targets and the most dominant clones were selected for downstream characterizations (Fig. 2a). Because the *K*_*D*_s of this first-generation *TRACeRs* landed in high nM range (Supplementary Fig. S6), we conducted one round of error-prone PCR and identified mutations that improve affinity without sacrificing specificity. Three key mutations identified were embedded into the original selected *TRACeRs* (Supplementary Fig. S7). This process improved their affinity by one-to two-orders of magnitude with minimum cross-reactivity (Fig. 2b).

**Figure 2.**
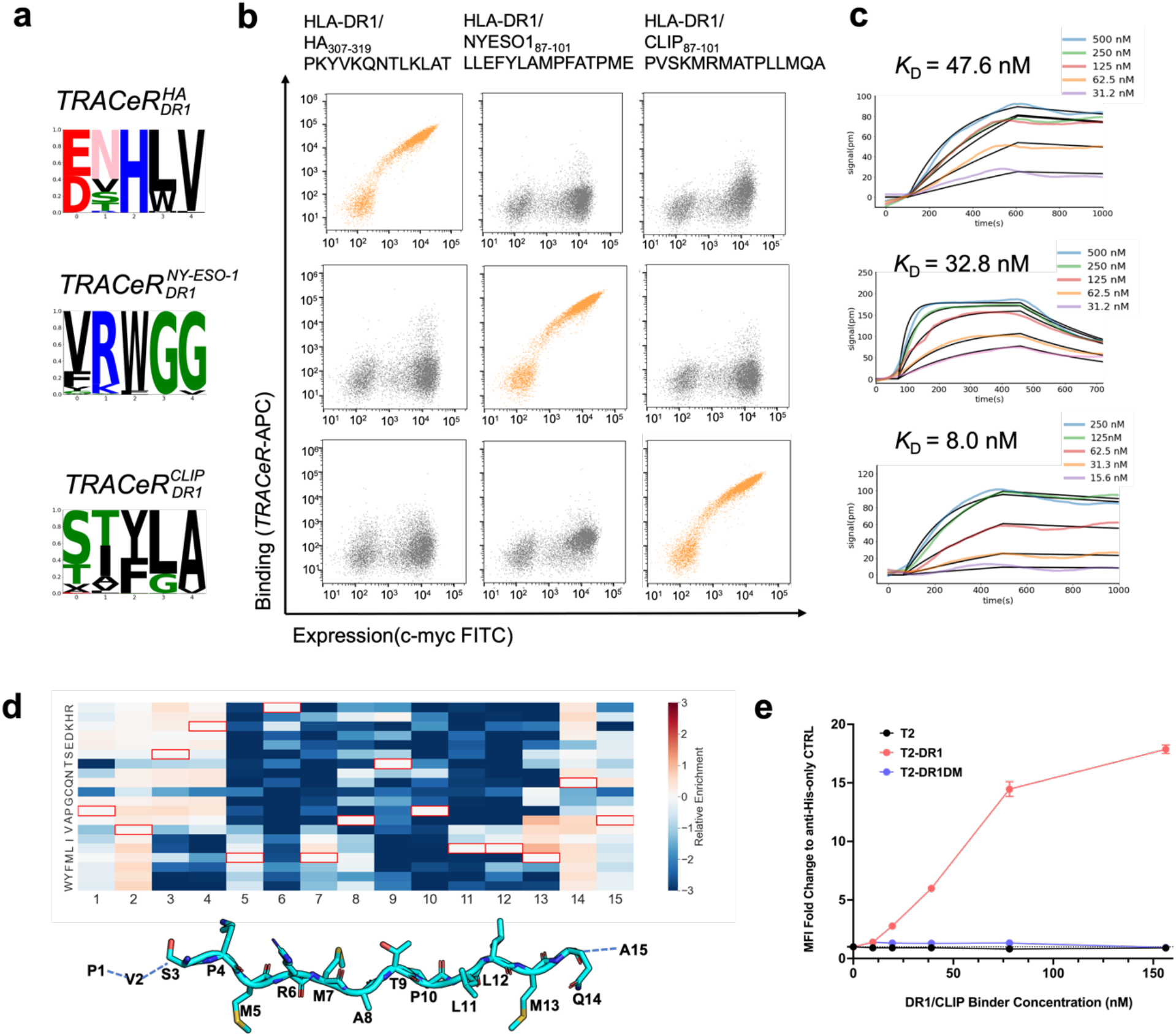
Specificity determination and biophysical characterization of *TRACeR* molecules. **a**. Sequence convergence of the ARE from the final sorting round. The most enriched clone for each target from the final sorting round was carried over for biophysical characterization and specificity determination. **b**. Target specificity determined with yeast surface display. Antigen-focused pMHC-II binders showed high specificity for their cognate antigens and minimal cross-reactivity with other targets. Staining concentration: on target, 25 nM tetramer (100 nM MHC + avidity); off target, 100 nM tetramer (400 nM MHC + avidity). **c**. Binding kinetics of *TRACeR* molecules determined by SPR. Fitting data are summarized in Supplementary Table 2. **d**. Relative enrichment heatmap of Site Saturation Mutagenesis (SSM) data on the CLIP peptide stained by 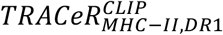. Relative enrichments were calculated after 3 rounds of FACS. Enrichment (red) indicates higher affinity, depletion (blue) indicates lower affinity. Boxed cells depict wildtype identity. The structural figure shows the side chain orientations of the antigen. **e**. The binding signal quantification relative to labeled anti-His background. Soluble 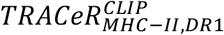 were used to stain T2 cells transfected with: non-transfected (DR1-, CLIP-, black), DR1 (DR1+, CLIP+, red), DR1DM (DR1+, CLIP-, blue). (n=3 biological replicates)

All identified *TRACeRs* were recombinantly expressible in the *E. coli* BL21(DE3) strain as soluble proteins (Supplementary Fig. S8). We collected the monomeric peak from size exclusion chromatography (SEC) and characterized binding affinity with surface plasmon resonance (SPR). All binding affinities landed in the low nM range, which is suitable for most applications^28^ (Fig. 2c, Supplementary Table 2). *TRACeRs’* specificity was further investigated with SPR using all the available targets, and no off-target activity was observed with concentration up to 2 μM, confirming the specificity pattern demonstrated with yeast surface display (Supplementary Fig. S9). Circular Dichroism spectra revealed distinct helical structures with melting transitions occurring at 60-80 °C, demonstrating high thermal stability (Supplementary Fig. S8). The solubility, stability, and ease of production of *TRACeRs* collectively establish it as a robust tool for addressing scientific inquiries in MHC-II biology and for exploring therapeutic applications.

To further demonstrate the specificity of binders developed with the *TRACeR-II* platform, we used yeast surface display to express a Site Saturation Mutagenesis (SSM) library with all 15 positions of the CLIP_87-101_ antigen mutated to all 20 amino acids, one at a time, on HLA-DR1^29, 30^ (Supplementary Fig. S10). The yeast library was stained with 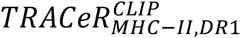 to enrich for the binding population. The breadth of binding specificity was revealed by calculating the enrichment of each amino acid at each position relative to the wild type (Fig. 2d)^31,32^. The CLIP antigen’s positions 5-13 are conventionally called the core region which sits within the MHC-II antigen binding cleft, and the other residues overhang the open binding pocket. The overhangs of the peptide outside the groove exhibit tolerance to multiple amino acid identities, aligning with the expectation that neither MHC-II scaffold nor 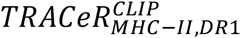 is anticipated to interact with these residues. In the core region, the combined influence of 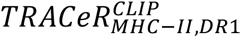 selectivity and MHC-II binding restriction renders the recognition highly specific. Positions 5, 8, 10, and 13 are the anchoring positions^16^ where the sidechains are oriented towards the bottom of the MHC-II peptide binding cleft, therefore their specificity are controlled by the MHC. Positions 6, 7, 9, 11, 12, are fully or partially solvent exposed and accessible to the ARE. Binding of *TRACeR* is sensitive to mutations at the following positions: Position 6 only tolerates K and R; Position 7, M, L, I; Position 9, S, T, H; Position 11 and Position 12, I, L, V. Overall, our mutagenesis analysis demonstrates the exquisite specificity rendered by the single ARE loop.

In addition, we investigated *TRACeR’s* ability to bind endogenously processed CLIP peptides presented on T2 cells. CLIP antigen is derived from the Class-II-associated invariant chain, and it is naturally required for folding for all MHC-IIs (Supplementary Fig. S11a). We used *TRACeRs* to stain the T2 cells transfected with HLA-DR1, HLA-DR1 & HLA-DM and a negative non-transfected control in a flow cytometry experiment. The use of HLA-DM in the experiments was to generate an HLA-DR1+/CLIP– control, as the chaperone HLA-DM catalyzes the exchange of CLIP peptides with other higher affinity peptides^33^. CerCLIP.1 antibody served as a positive control to confirm CLIP surface expression^34^. As expected, 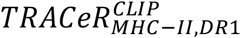 binds to cell surface HLA-DR1/CLIP in a concentration-dependent manner, while 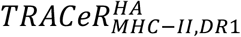 showed no binding signal at micromolar concentration (Fig. 2e, Supplementary Fig. S11b, c). Presence of HLA-DM abolished binding of 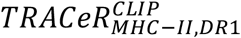, highlighting the peptide dependence of *TRACeR* binding to MHC. Our results validate that the *TRACeR* has the ability to bind cellularly-processed antigens and does not exhibit cross-reactivity with random cellular peptides introduced through DM-catalyzed peptide exchange.

### Multi-allelic compatibility with high sensitivity to peptide conformation

The unique binding mode of *TRACeR-II* platform displays fundamental differences from those of TCR and antibody. Structurally, MAM interacts with the MHC peptide-binding groove on the α chain, which is shared among different HLA-DR molecules. *TRACeR-II* platform inherits the property of MAM to bind different alleles, which can greatly simplify the development of antigen-focused binders capable of targeting polymorphic MHC-IIs^24^. To test these assumptions, we first investigated 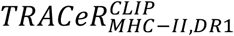’s ability to bind to different HLA alleles presenting the same CLIP peptide. 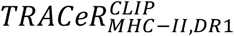 was found to also bind HLA-DR4/CLIP (Fig. 3a) with similar affinity and specificity pattern of HLA-DR1 (Fig. 3b and Supplementary Fig. S12). Surprisingly, we observed no binding signal with HLA-DR3/CLIP or HLA-DR15/CLIP. Comparing the CLIP antigen conformations in the available x-ray structures of HLA-DR1 (PDB ID: 3QXA) and HLA-DR3 (PDB ID: 1A6A) revealed a subtle structural difference in the threonine at position 9 (Supplementary Fig. 13). This result suggests that the *TRACeR-II* ARE loop is highly sensitive to the difference in peptide conformation^35^. To further investigate the compatibility of *TRACeR-II* to HLA-DR3 and HLA-DR15 alleles, we created peptide-specific binders targeting HLA-DR3 with human HER2/neu_389-403_ (SANIQEFAGCKKIFG)^36^ and HLA-DR15 with αSynuclein_32-46_ (KTKEGVLYVGSKTKE)^37^ (Fig. 3c, d and Supplementary Fig. 14, 15). We found that *TRACeR-II* is compatible with these targets, and therefore 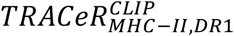 not binding CLIP peptide on these alleles is because of the antigen conformation.

**Figure 3.**
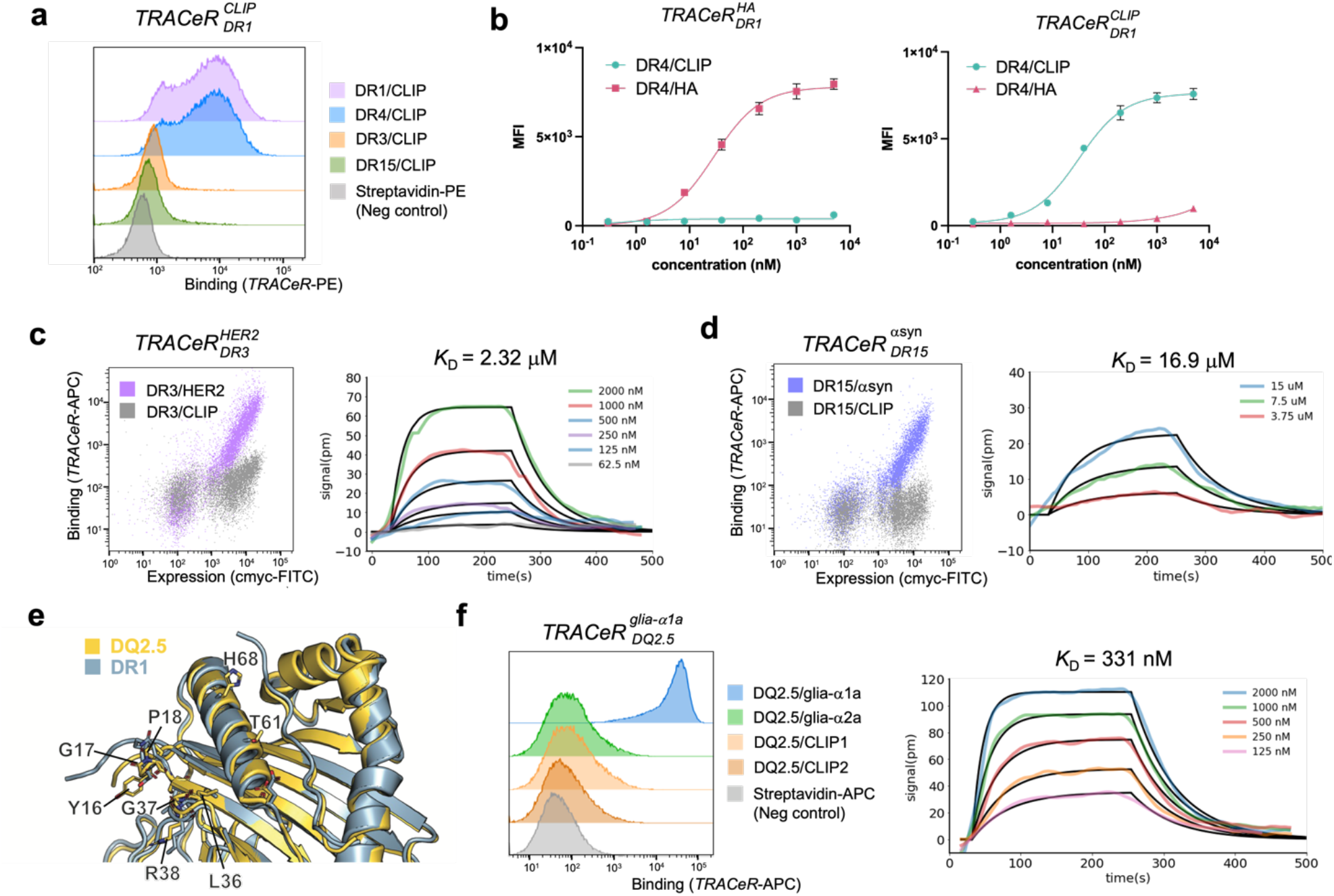
Muti-allelic compatibility of *TRACeR* platform. **a**. Histogram of 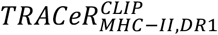 binding the CLIP peptide frame (87-101) presented by different HLA-DR alleles. 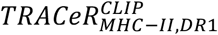 is displayed on the yeast surface and stained with 50 nM pMHC tetramer. **b**. Titration of HLA-DR4 MHC tetramer on yeast displaying 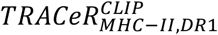 (right) or 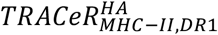(left) (n=3 technical replicates). **c**. Left: 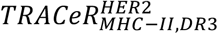 displayed on the yeast surface stained with HLA-DR3/HER2 (positive control) and HLA-DR3/CLIP (negative control). Right: Binding kinetics of 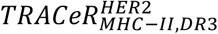 with HLA-DR3/HER2 determined by SPR. **d**. Left: 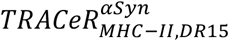 displayed on the yeast surface stained with HLA-DR15/αSyn (positive control) and HLA-DR15/CLIP (negative control). Right: Binding kinetics of 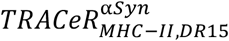 with HLA-DR15/αSyn determined by SPR. **e**. Positions on HLA-DQ2.5 allele α chain which do not share the same side chain identity with DRA1*01:01. Only residues within 5 Å from *TRACeR* are shown. **f**. Left: 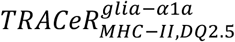 displayed on the yeast surface and stained with 50 nM pMHC tetramer with different antigens presented on HLA-DQ allele. Right: Binding kinetics of 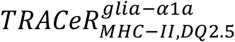 with HLA-DQ2.5/glia-α1a determined by SPR.

Beyond HLA-DR alleles, we additionally tested *TRACeR-II*’s compatibility with an HLA-DQ allele, which does not share the same α chain (Fig. 3e). We were able to develop a binder for a celiac disease antigen on the DQ allele, specifically HLA-DQ2.5 (HLA-DQA1*0501 and HLA-DQB1*0201) with the glia-α1 epitope QLQPFPQPELPY (Fig. 3f and Supplementary Fig. 14, 15), demonstrating that *TRACeR-II* is a versatile platform compatible with diverse MHC backgrounds. Given the high sensitivity to intricate peptide conformation, we would expect that *TRACeR*s can resolve multi-frame loading of antigens. HLA-DQ2.5 allele can present CLIP peptide in two distinct frames when loaded with CLIP1(84-101) and CLIP2(92-107)^38^. CLIP1’s loading frame on HLA-DQ2.5 is equivalent to that on HLA-DR1. In a tetramer staining experiment on yeast surface display, we found that 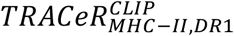 can recognize only the CLIP1 frame on DQ2.5, but not the CLIP2 frame (Supplementary Fig. 16). This further supports the selective nature of *TRACeR-II* binders.

In summary, the *TRACeR-II* platform emerges as a versatile, multi-allelic tool with a unique binding mode that encodes peptide specificity, offering a potential solution to simplify the challenges of polymorphism in developing MHC-II binders.

### Structural validation by cryo-electron microscopy

To validate our engineering hypotheses and gain insight into the antigen recognition mechanism, we determined the cryoEM structure of 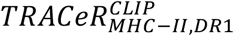 -DR1/CLIP complex at 2.34 Å resolution (Fig. 4a, Supplementary Fig. S17). The c44H10 Fab was co-purified with the *TRACeR*-MHC complex to facilitate particle alignment and achieve high resolution^39^.

**Figure 4.**
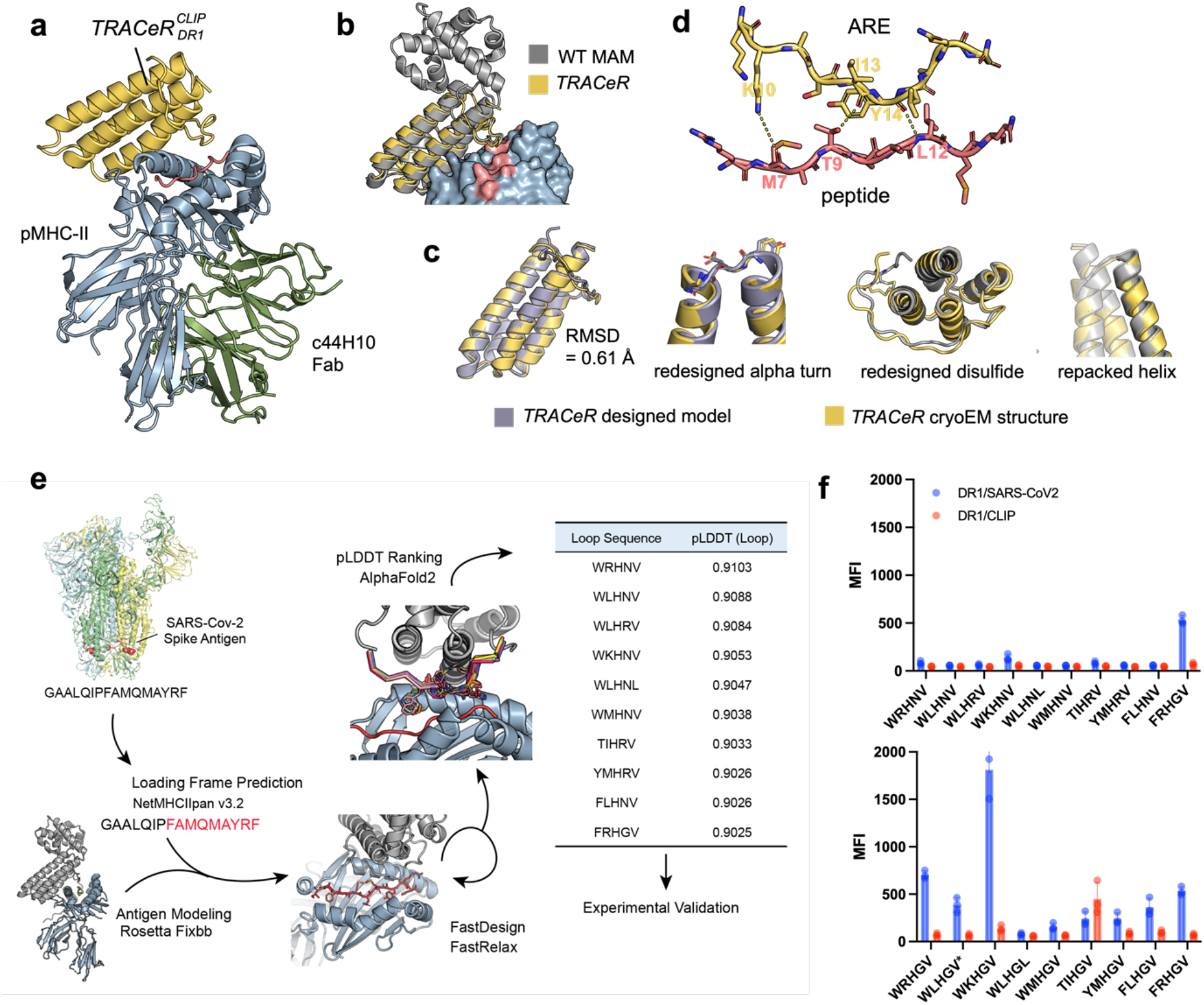
CryoEM structure reveals a unique binding mode. **a**. CryoEM structure of 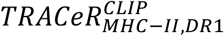⋅ HLA-DR1/CLIP ⋅ c44H10 Fab ternary complex. **b**. Superposition of the wildtype MAM on the *TRACeR* cryoEM structure in **a. c**. Comparison of the designed *TRACeR* model to the cryoEM structure. left to right: global superposition; redesigned alpha turn; designed disulfide; repacked helix. **d**. Zoomed-in view of the ARE-peptide interface. **e**. Computational design scheme for 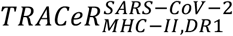, detailed method description and design scripts in Supplementary Appendix 1. **f**. Experimental validation of designed *TRACeRs* displayed on the yeast surface, stained with 50 nM HLA-DR1/SARS-CoV-2 and HLA-DR1/CLIP tetramers. (n=3 technical replicates). *Both WLHNV and WLHRV turned into WLHGV when mutating the 4^th^ residue to G.

As expected, the ternary complex structure reveals that the binding mode of 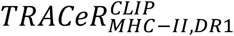 closely resembles that of MAM and MHC-II (Fig. 4b). The structure of 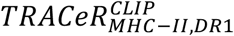 shows a remarkable agreement with the computationally designed model in Figure 1c with a Cα RMSD of 0.61 Å. The redesigned alpha-alpha turn of *TRACeR* aligns perfectly with the experimentally determined structure. The designed disulfide holds the ARE in place as designed, although Rosetta had selected a different rotamer on one of the cysteine residues (Fig. 4c).

For the ARE-antigen interface, we observed that the ARE interacts with the peptide using both polar interactions and side chain-derived shape complementarity. As for polar interactions, Lys10_TRACeR_ forms a water-mediated hydrogen bond with the carbonyl group of Met7_CLIP_; the carbonyl group of Ile13_TRACeR_ with Thr9_CLIP_; the carbonyl of Tyr14_TRACeR_ with amide of Leu12_CLIP_ (Fig. 4d).

The binding selectivity was surprisingly achieved with a small binding footprint on the peptide. Given that the limited polar interactions observed did not adequately account for the binding specificity, we postulate that antigen recognition is primarily accomplished by a passive recognition mechanism. To successfully engage with a specific peptide, *TRACeR* relies on a stringent compatibility criterion for all side chains involved, ensuring the absence of any steric conflicts. Any clashes or overlaps between molecular components will lead to a depletion of the binding signal. These structural features also further verify the specificity pattern that was observed in Figure 2d. For every amino acid on the antigen within the reach of the *TRACeR* ARE loop (positions 6, 7, 9, 11, 12), their specificities are clearly explained by the cryoEM structure. Mutations in the close proximity range (positions 9-12_CLIP_) are highly restricted because the ARE loop exerts dominant influence over the tolerated amino acid identities; in this region, only amino acids with similar size and property are allowed. The flanking positions (positions 5, 6, 7, 8 and 13) showed mutational preference to a subset of amino acids upon *TRACeR* binding, and this is likely due to the loading conformation of the antigen backbone in the MHC. The structure also supports the critical change in the conformation of Thr9_CLIP_ that abolishes HLA-DR3 binding. The hydrogen bond between Thr9_CLIP_ and the carbonyl of Ile13_TRACeR_ likely contributed to the conformational specificity (Supplementary Fig. S13).

### A unique antigen recognition mechanism enabling direct structural design

The agreement between the cryoEM structure and the original MAM complex suggests that the binding mode for *TRACeR-II* is indeed constant. This opens the possibility that specific MHC-II antigen binding can be directly designed into this very constrained conformational space. The simplistic loop-on-loop interaction reduced the modeling problem to a tractable scale. We tested this notion with an antigen of unknown complex structure (a SARS-CoV-2 antigen on HLA-DR1 with the sequence GAALQIPFAMQMAYRF). (Fig. 4e).

We incorporated Rosetta modeling and AlphaFold2 prediction tools to attempt the design challenge. The protocol is presented in Figure 4e. We first predicted the tightest antigen binding frame to the MHC-II with the NetMHCIIpan online server^40^. Residues were then threaded onto the MAM-DR1 complex structure (PDB ID: 2ICW) following the predicted frame. We used Rosetta to model the ARE and its interactions with the antigen. We forced the same cysteine residues on the *TRACeR* scaffold and relaxed the environment near the disulfide. During the design process, the same five residues used in the original library were allowed to mutate. These five residues and all the neighboring residues (including MHC, antigen and *TRACeR*) within 12 Å were allowed to move during the FastDesign and FastRelax steps (Supplementary appendix 1 and 2). The resulting models were then scored by Alphafold2 and ranked by their pLDDT scores^41^. The top 10 designs were tested with yeast surface display, and one of the sequences showed binding to the designated target (Fig. 4f, top). All the models scored similarly by pLDDT, and the ARE sequence that showed binding (FRHGV) was not among the best-scored models ranked by Rosetta metrics. We inspected the differences between the design models and noticed a positive ϕ backbone angle at the 4^th^ position in the ARE loop, where a glycine was designed onto the successful sequence while other designs have polar residues (Asn or Arg). A strict design rule dictates that glycine should have been used for this position, and the design of polar residues is likely a Rosetta artifact. We then tested variants of the unsuccessful sequences with glycines in their 4^th^ position to account for the Rosetta artifact. We rescued two more antigen-specific designs with better affinity (Fig. 4f, bottom). ARE sequence that yielded the highest binding signal on yeast was purified as a recombinant protein and has an SPR-measured *K*_D_ of 3.78 μM (Supplementary Fig. S18). Interestingly, although no specific negative design was carried out, the designed loop showed specificity against an irrelevant peptide (CLIP). That might be attributed to the strict requirement of compatibility at the ARE region as suggested by the structural analysis.

The results showed that the restricted sampling space of the *TRACeR* binding loop enables the direct design by simple physical based rotamer and loop conformation sampling. Utilizing the design scheme reduces the experimental test scale from 10^6^ to the order of 10s. However, the Rosetta artifacts affected the success rate and more advanced design methods might place the glycine correctly. Furthermore, the design relies on the correct prediction of the peptide frame, so we would anticipate better success rate on pMHCs with solved complex structures.

## Discussion

MHC-IIs play a central role in the adaptive immunity, as they are responsible for antigen presentation that leads to CD4+ T cell activation and differentiation. As a critical hub for antigen information exchange, MHC-IIs are strongly implicated in human diseases, from autoimmune diseases to cancer, and thus the characterization and discovery of MHC-II-related antigens are fundamental for the studies of pathogenesis and potential theragnostic solutions. The demand for antigen-specific MHC-II binding proteins is high, but despite the immense progress with TCRs and TCR-like antibodies, the labor-intensive engineering processes have not met the theragnostic needs. We discovered that the unique binding features of the superantigen MAM could potentially offer the simplest solution by reducing the antigen recognition interface to a single loop. Remarkably, a single loop tethered on one end with a disulfide bond could achieve discrimination of single mutations on MHC-II presented peptides. It can encode peptide selectivity with as few as 5 residues in the ARE, and this region can be expanded in case additional interactions with the peptide are needed.

*TRACeR-II*’s single loop-based interface inspired us to carry out computational designs using only the antigen’s sequence. As a proof-of-concept, we successfully designed a *TRACeR* against an arbitrarily chosen antigen. A design pipeline like this could potentially be automated to address the expanding antigen repertoire. The success with this effort could vastly reduce the amount of labor and time required for developing binders for newly discovered pMHC-II antigens.

The crystal structure of the ternary complex of MAM, TCR, and MHC-II has been available in the PDB since 2007. As a superantigen, MAM presents several unique features: it is the only known naturally occurring superantigen that binds near the antigen; it interacts with the α chain of MHC-II and can bind not only different HLAs, but also mice MHCs. Wildtype MAM is not known to be antigen-specific and has not been explored in that regard. However, through protein design, we were able to build from MAM a highly stable, versatile, and designable MHC-II binding platform. We have demonstrated that peptide-specific binders for multiple HLA alleles, including HLA-DR1, DR3, DR4, DR15 and DQ2.5, can be selected from a single “peptide-focused” ARE library. Our results presented an alternative strategy for addressing the challenge of MHC-II peptide targeting, providing a promising foundation for developing potent tools to explore MHC-II biology and applications in cancer, viral infection, and autoimmune diseases.

## Acknowledgement

We thank Alexander Muselman and Dr. Edgar Engleman for help and support on animal experiment. We thank Christian A. Choe for helpful discussion and Dr. Nadia Roan for helpful comments and feedback on the manuscript. We thank the NIH Tetramer Core Facility (contract number 75N93020D00005) for providing all HLA-DR tetramer reagents. We thank Rui Yan at the Janelia CryoEM facility and Elizabeth Montabana at the Stanford University Cryo-Electron Microscopy Center for help in microscope operation and data collection. H.D. is supported by Stanford Bio-X Graduate Fellowship; J.L. by Stanford Graduate Fellowship Award; B.B. by NIH Biotechnology Training Program; Y.L. by Stanford Maternal and Child Health Research Institute; A.K. is supported by an Ontario Graduate Scholarship (OGS); J.-P.J. is supported by the Ontario Early Researcher Award program and the Canada Research Chairs program. E.D.M. is supported by NIH (R01AI159260) and the Stanford Maternal and Child Health Research Institute; K.C.G. is supported by Howard Hughes Medical Institute and NIH (5R01AI103867). P.-S.H. is supported by NIH (R01GM147893), American Cancer Society (ACS 134055-IRG-218), BASF CARA project, and Discovery Innovation Fund.

## Author contribution

Project Conception: H.D., J.L., P.-S.H.

Manuscript writing: H.D., J.L., P.-S.H.

Computational Modeling: J.L., H.D.

*TRACeR* experimental development and characterization: H.D., J.L., B.B., H.Y., Y.L., A.M., U.P. Cryo-EM and structural determination: K.M.J., X.Y., K.C.G.

Development and production of c44H10: A.K., J.-P.J.

Immunogenicity assays: H.D, R.A.P.S.

Cell assays: Y.L., E.D.M.

Contribution to data, data analysis, experimental design and manuscript feedback: all authors.

## Methods

### Library design and production

*TRACeR* library with NNK degenerate codon at the 5 loop positions was ordered from IDT as Ultramer™. The Ultramer™ are then assembled and amplified.

For library transformation, *Saccharomyces cerevisiae* yeast EBY100 cells were transformed with insert DNA and linearized pCTCON2 plasmid using the established protocol^42^. After transformation, cells were grown overnight in SDCAA media at 30 °C, passaged once, and stored in 20% glycerol solution at −80 °C.

### Yeast display and library screening

Transformed yeast cells were grown in SDCAA media. For induction of expression, yeast cells were centrifuged at 2000 × g for 5 min and resuspended in SGCAA media supplemented with 0.2% glucose at the cell density of 1e7 cells per ml and induced at 30 °C for 16-24 h. Cells were washed with PBSA (phosphate buffer saline with 0.5% BSA) and labeled with pMHC tetramer or monomer, together with anti-c-myc fluorescein isothiocyanate (FITC, Miltenyi Biotec). After incubation for ∼1 h at room temperature, cells were washed twice and resuspended in PBSA, then run on a Sony SH800 cell sorter.

### Next Generation Sequencing

Up to 1 × 10^8^ yeast cells from a growing culture were taken to perform a zymoprep using the ZymoprepII kit (D2004, Zymo Research). The *TRACeR* sequences were amplified from the extracted plasmids and the adapter sequences were added by PCR. The PCR product was gel purified and sent for NGS (Amplicon-EZ, genewiz). The most enriched clones were taken for subsequent experimental characterization.

### Error-prone PCR and affinity maturation

The codon-optimized original CLIP binder gene was ordered from Twist Bioscience. Error-prone PCR reactions were carried out following the protocol provided by GeneMorph II Random Mutagenesis Kit (200550, Agilent). Primers were designed to flank residue 1 to 119 of the binder. For each 50 μL reaction, 10 ng of template (as calculated by the amount of CLIP binder gene) was used. The reactions were then gel purified. 2 μg of insert was transformed into competent EBY100 cells with linearized pCTCON2 vectors. The transformed library size was 1.2e7. 3 rounds of cell sorting were carried out with cells co-stained by both CLIP tetramer and HA tetramer. DNA from both the original library and the resulting population were extracted and sent for NGS. The enrichment of mutations was analyzed using python. Enriched point mutations were combined for subsequent binding affinity tests, and the combination that produced the highest binding affinity was selected.

### Protein expression and purification

Genes encoding the designed protein sequence were synthesized and cloned into pET-24a(+) *E*.*coli* plasmid expression vectors (Genscript, C-terminal 6X His tag). Plasmids were then transformed into chemically competent BL21(DE3) *E. coli* (ZYMO research). The cells were cultured in 2xYT media at 37 °C until OD reached 0.6∼0.8. Protein expression was then induced with 1 mM of isopropyl β-D-thiogalactopyranoside (IPTG) at 16 °C. After overnight expression, cells were collected and resuspended with 50 mM Tris buffer (pH = 8.0, 300 mM NaCl) and frozen at −80 °C until purification. The cell pellet was thawed, sonicated and purified by nickel affinity followed by size exclusion chromatography (Superdex™ 75 10/300GL or Superdex™ 75 10/300GL Increase GE Healthcare). All protein samples were characterized by SDS-PAGE. Protein concentrations were determined by absorbance at 280 nm measured with a Nanodrop spectrophotometer (Thermo Scientific) using predicted extinction coefficients.

### Circular Dichroism

CD spectra were measured on a JASCO CD spectrophotometer in a 1-mm pathlength cuvette (Hellma). Protein samples were at ∼0.2 mg/mL in the 50 mM Tris buffer. Temperature melts were from 20 to 95 °C. and CD signal at 222 nm were monitored in 1 °C increments per minute, with 10 s of equilibration time and 1 s digital integration time. Wavelength scans (200-260 nm) were collected at 20 °C and 95 °C, and again at 20 °C after fast refolding.

### Surface Plasmon Resonance

The SPR experiments were performed on OpenSPR (Nicoya). Biotinylated pMHC monomers were immobilized on streptavidin coated sensor using biotin-streptavidin sensor kit (SEN-AU-100-10-STRP-KIT). pMHC was immobilized at a density of 1500-2000 RU on channel 2. Channel 1 was left blank to serve as a reference surface. To collect binding data, *TRACeRs* in PBS + 0.05% Tween-20 were injected at a concentration series at room temperature. Association and dissociation time was adjusted based on a pilot run on each sample for best fitting results (association time is 300 s by default but was extended to up to 550 s for samples with lower on-rate). Raw data was processed with Savitzky-Golay filtering for removing high dimensional noise^43^. Kinetic parameters were acquired by fitting to a 1:1 interaction model using the TraceDrawer software (Nicoya). Fitting was performed with raw data before Savitzky-Golay filtering.

### Immunogenicity assay

Animal studies were approved by the Stanford Administrative Panel on Laboratory Animal Care (APLAC). BALB/c mice (female, 6-8 weeks old, n=5 per group) were randomly separated into groups, anesthetized and dosed (retro orbital injection) with Tris buffer, 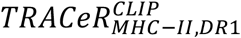 (in 50 mM Tris buffer, pH=8) or monoclonal antibodies mIgG (Innovative IR-MSBC-GF) or hIgG (Innovative IR-HU-GF-ED), with a dose of 0.2 mg/kg. Injections were performed every two days for two weeks at day0, day2, day4, day6, day8, day10, and day12. Blood was collected at day 0, day7 and day14 by retro orbital bleed using micro-hematocrit capillary tubes (Fisher). Mice were weighted before every injection. Serum was separated by centrifuging the blood samples in PCR tubes. For mouse experiments, researchers were not blinded to animal identity.

### ELISA

ELISA was used to evaluate the levels of *TRACeR*, mIgG, hIgG, and BSA-specific IgG antibodies in mice serum. Maxisorp plates from Thermo Scientific-Nunc were coated with 100 ng per well of *TRACeR*, mIgG (Innovative IR-MSBC-GF), hIgG (Innovative IR-HU-GF-ED), or BSA (LAMPIRE Biological laboratories, cat no. 7500804) in PBS, and incubate overnight at 4 °C. The plates were then blocked with 5% nonfat milk powder in PBS for 1 h at room temperature, followed by three washes with wash buffer (PBS-T: phosphate-buffered saline containing 0.05% Tween 20). The samples were diluted in a buffer consisting of 1% nonfat milk powder in PBS-T, added to the wells, and incubated for 1 h at room temperature. After three washes with PBS-T, the plates were incubated with horseradish-peroxidase conjugated goat anti-mouse IgG (diluted 1/5,000) secondary antibodies (ThermoFisher 62-6520) for 1 h at room temperature. Following five washes with PBS-T, TMB substrate (KPL 52-00-03) was added to the wells and incubated for 30 min at room temperature. The development of color was stopped by adding 50 μL of 1 M HCl, and the plates were read at 450 nm to measure the relative optical densities. The average optical density of blank wells was subtracted from the readings to calculate the reported values.

### Specificity Assessment of 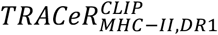with yeast surface display

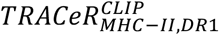with avi-tag was ordered from genscript in pET24-a(+) vector. The biotinylated protein was made using BL21 strain containing BirA plasmid (BLB-201, Amid Bioscience) following the product protocol. The yeast display construct ptDR1 was kindly provided by Dr. Mellins ^29, 30^. The site saturation mutagenesis library of CLIP 87-101 was ordered from Twist Bioscience and PCR amplified. The insert library was transformed with the double digested ptDR1 vector into EBY100 yeast strain by electroporation. The yeast culture was grown in SDCAA media and induced at OD 2-5 in SGCAA media for 18 hours at 30 °C. The induced culture was stained with anti-HA-APC (cat. 901523, BioLegend) and tetramerized streptavidin-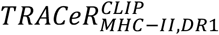-PE. The double positive population was enriched over 3 rounds. Gene extracted from the unselected population and the final round product was sent for NGS. Data was analyzed using python by calculating the relative enrichment of each amino acid identity at each peptide position.

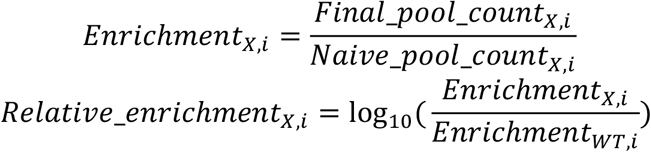

### Construction of HLA-specific T2 cell lines

To test the binding affinity and specificity of *TRACeR* with the naturally processed pMHC-II complexes, cell lines expressing single HLA-DR1 allele were constructed by stable transfection of HLA-DR1, or stable co-transfection of HLA-DR1/HLA-DM (DMA and DMB). T2 is a class II-deficient (HLA-II-DM-DO-) TxB hybrid cell^44, 45^. HLA-DM catalyzes CLIP peptide dissociation from HLA-DR molecules and facilitates loading of other peptides; this function is critical for exchange of CLIP, particularly for alleles with high CLIP affinity such as HLA-DR1^33, 46^. Thus, at the cell surface, T2-DR1 transfectants express CLIP-associated DR1 complexes, while T2-DR1DM transfectants express primarily HLA-DR1 complexes with other peptides. T2 and its transfectants were cultured in IMDM, GlutaMAX supplemented media (Thermo Fisher Scientific) with 10% HI FBS, 1% penicillin/streptomycin (P/S) and 0.5 mg/ml G418 (to maintain selective pressure on DR transfectants) at 37 °C with 5% CO_2_. Expression of the HLA-DR protein was detected by anti-HLA-DR (clone L243) and CLIP-associated HLA-DR complexes was detected by anti-CLIP clone (cerCLIP.1). Expression was evaluated by flow cytometry before performing *TRACeR* binding experiments.

### *TRACeR* binding test to T2 cell lines

Sufficient cells (1 × 10^6^ per tube) from the cell culture were collected, washed with 4 °C PBS, resuspended in 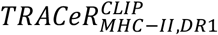 containing PBS buffer, and incubated at room temperature for 1 h. After incubation, cells were washed with FACS buffer to remove excessive unbound *TRACeR*, followed by the staining with anti-His-APC (1:20, clone J095G46, BioLegend cat# 362605) with Fc block (1:20, BioLegend cat# 411302) at 4 °C for 40 min. Stained cells were washed with FACS buffer and then resuspended with 200 μL FACS buffer per sample containing propidium iodide (PI) dye (40 ng/test, Invitrogen™ cat# 00699050) to indicate cell viability. To get binding kinetics to the cell surface presenting HLA-II/CLIP complex, a titrated amount of 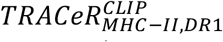(0, 9.77, 19.53, 39.06, 78.13, 156.25 nM) was tested. 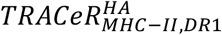 binder was used as a control for non-specific binding. *TRACeR* binding signal was defined as the mean fluorescence intensity (MFI) of the anti-His channel of the viable (PI-) singlet population. Flow cytometry data were acquired using a BD FACSymphony™ A5 Cell Analyzer (BD Biosciences) in Stanford Shared FACS Facility (SSFF) and analyzed using FlowJo software (version 10.8.1).

### Production of c44H10 Fab

FreeStyle 293-F cells (Thermo Fisher Scientific) were split to a density of 0.8 × 10^6^ cells/mL 1 h before transfection. Cells were transfected using FectoPRO Reagent (Polyplus) following manufacturer instructions at a 1:1 DNA to FectoPRO ratio. 90 μg of plasmid DNA was used for transfection (2:1 ratio of heavy and light chain DNA plasmids) for every 200 mL of cell culture. Transfected cells were incubated in a 37°C, 5% CO_2_ shaking incubator for 5 to 7 days to allow for the expression and pairing of heavy and light chain gene products. Transfected cell culture supernatants were collected and filtered through 0.22 μm Steritop filters (Millipore Sigma). Recombinant c44H10 Fab was purified by KappaSelect affinity chromatography with 100 mM glycine, pH 2.2 elution and immediate 1 M Tris, pH 9.0 neutralization, followed by MonoS ion exchange chromatography using 20 mM NaOAc, pH 5.6 + 1 M KCl.

### Production of HLA-DR1/CLIP for cryoEM study

The expression construct for HLA-DR1/CLIP was described previously^47^. In brief, residues 1-182 of the DR α chain were fused with an acidic zipper and cloned into a pD649 vector. The CLIP peptide was fused with residues 1-190 of DRβ chain via a flexible linker containing thrombin cleavage site. The CLIP-DR1 β construct was fused with a basic zipper and cloned into a pD649 vector. To express CLIP-HLA-DR1, 15 μg plasmid of α chain and 15 μg plasmid of β chain were co-transfected into 30 mL of Expi 293 cells at a density of 2.5 × 10^6^ cells/ml and expression proceeded for 3 days. The supernatant contained CLIP-HLA-DR1 was sequentially purified by Nickel NTA and a Superdex™ 200 increase column. The fractions containing CLIP/DR1 are pooled, and quality of protein was assessed by SDS-PAGE gel. To prepare the *TRACeR* ⋅ HLA-DR1/CLIP complex, the HLA-DR1/CLIP was incubated with 3C protease to remove the acid-basic zippers.

### *TRACeR* ⋅ HLA-DR1/CLIP ⋅ Fab complex purification

10.6 nmoles of 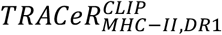 and 6.4 nmoles of c44H10 Fab were added to 5.3 nmoles of HLA-DR1/CLIP. After incubating for 30 min on ice, the protein complex was purified by FPLC on a Superdex™ 200 increase column. Selected fractions were analyzed by SDS-PAGE and concentrated to 2 mg/ml.

### CryoEM microscopy

The protein complex was diluted to 0.5 mg/ml in the presence of 0.01% fluorinated octyl maltoside. Aliquots of 3 μl were applied to glow-discharged 300 mesh Quantifoil (1.2/1.3) grids, which were blotted for 2 s at 4°C and 100% humidity with an offset of −15 and plunged frozen into liquid ethane using a Vitrobot Mark IV (Thermo Fisher). Grids were screened at the Stanford cEMc on a 200 kV Glacios microscope (Thermo Fisher) equipped with a K3 camera (Gatan). A dataset was collected on a single grid using a 300 kV FEI Titan Krios microscope located at the HHMI Janelia Research Campus. Movies were collected using a 6eV wide energy filter at 165,000× magnification, corresponding to physical pixel size of 0.743 Å. Automated data collection was carried out using SerialEM with a total dose of 60 e^-^/Å^2^ in 42 frames over a nominal defocus range of −0.8 to −1.8 μm.

### Image Processing

All processing was performed in cryoSPARC^48^ (Supplementary Fig. S17). 21,328 movies were motion-corrected using patch motion correction. The contrast transfer functions (CTFs) of the aligned micrographs were determined using patch CTF and an initial stack of 100,993 particles was chosen from the first 704 micrographs by blob picking. After successive rounds of reference-free 2D class averaging, 2D classes corresponding to intact complexes were used as templates to pick 18,639,350 particles from the complete dataset. After 2D sorting to remove junk particles, a random set of 498,178 particles was used for ab initio reconstruction, followed by two rounds of 3D sorting of the full set of 15,580,106 non-junk particles. The resulting 3,706,646 particles were then local motion corrected and the volume was improved by non-uniform refinement. Due to mobility of the Fab constant domains, local refinement was performed using a mask to exclude these domains, and this 2.34 Å resolution map was used for subsequent real space refinement in phenix. Additional sharpening was performed with deepEMhancer^49^.

### Model Building and Refinement

Models of DR1 with peptide (PDB ID: 6R0E), 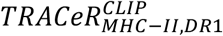, and the c44H10 variable domains (PDB ID: 8EUQ) were docked into the deepEMhancer map using ChimeraX^50, 51^. Sequences were corrected and the molecules were interactively rebuilt in Coot^52^ using the deepEMhancer map. Atomic coordinates and atomic displacement parameters were refined using phenix Real Space Refinement^53, 54^ using an autosharpened local refinement map, resulting in a final model with a Molprobity score of 1.61 and zero Ramachandran outliers^55^. CryoEM data collection, refinement, and validation statistics are presented in Supplementary Table 3. Model building and refinement software was installed and configured by sbgrid^56^.

### Data availability

Cryo-EM maps have been deposited in the Electron Microscopy Data Bank (EMDB) under accession code EMD-43499. Atomic coordinates have been deposited in the Protein Data Bank (PDB) under accession code 8VSJ.

