## Supplementary Information for "A general platform for targeting MHC-II antigens via a single loop"

**This PDF file includes**
Supplementary Figure S1-S17
Supplementary Table 1-3
Supplementary Appendix 1-2

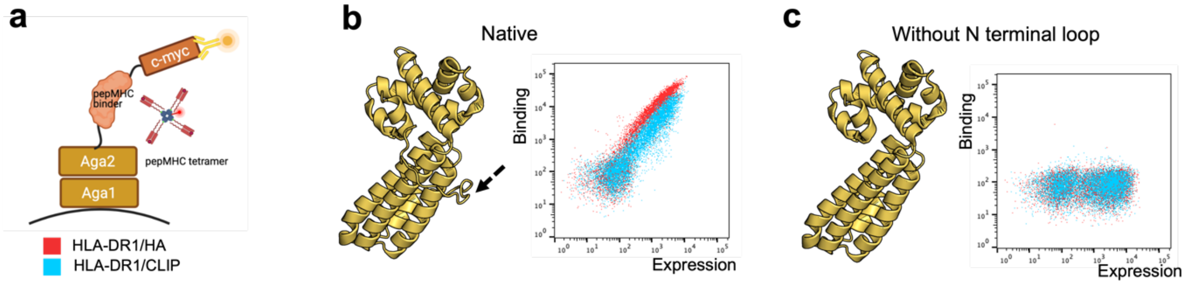


**Figure S1 The N-terminal loop (ARE region) is essential for antigen engagement**
Removal of the N-terminal loop from MAM completely abolishes binding to pMHC-II complexes. **a.** Schematic of the yeast surface display experiment. **B.** Native MAM displayed on the yeast surface and stained with HLA-DR1/HA tetramer (red) or HLA-DR1/CLIP tetramer (blue). The ARE region is labeled with the black arrow. **C.** MAM without residue 1-20 displayed on the yeast surface and stained with HLA-DR1/HA tetramer (red) or HLA-DR1/CLIP tetramer (blue). Staining concentration: 50 nM tetramers (200 nM pMHC + avidity).

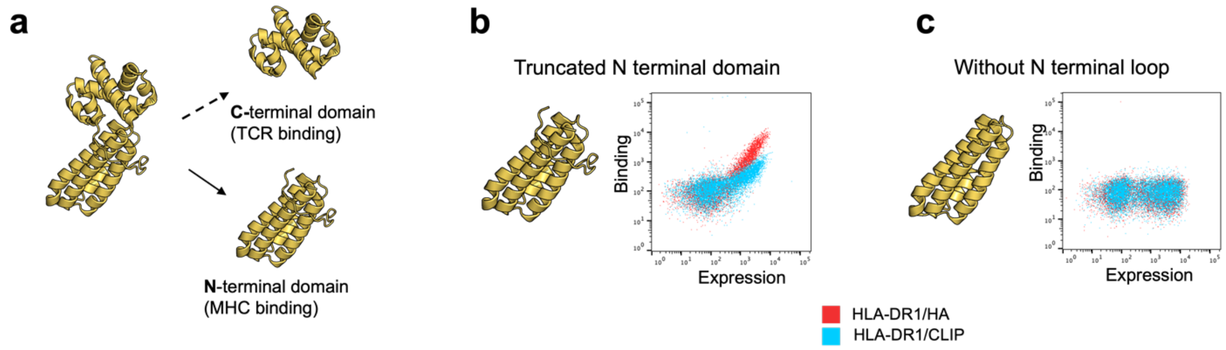


**Figure S2 MAM N-terminal domain maintains MHC binding**
**a.** Structure of MAM and domain splits. MAM is a two-domain protein that can be split into an N-terminal domain that binds MHC-II and a C-terminal domain that binds TCR. **b.** Truncated MAM N-terminal domain displayed on the yeast surface and stained with HLA-DR1/HA tetramer (red) or HLA-DR1/CLIP tetramer (blue). **c.** MAM N-terminal domain without residue 1-20 displayed on the yeast surface and stained with HLA-DR1/HA tetramer (red) or HLA-DR1/CLIP tetramer (blue). Staining concentration: 50 nM tetramers (200 nM pMHC + avidity).

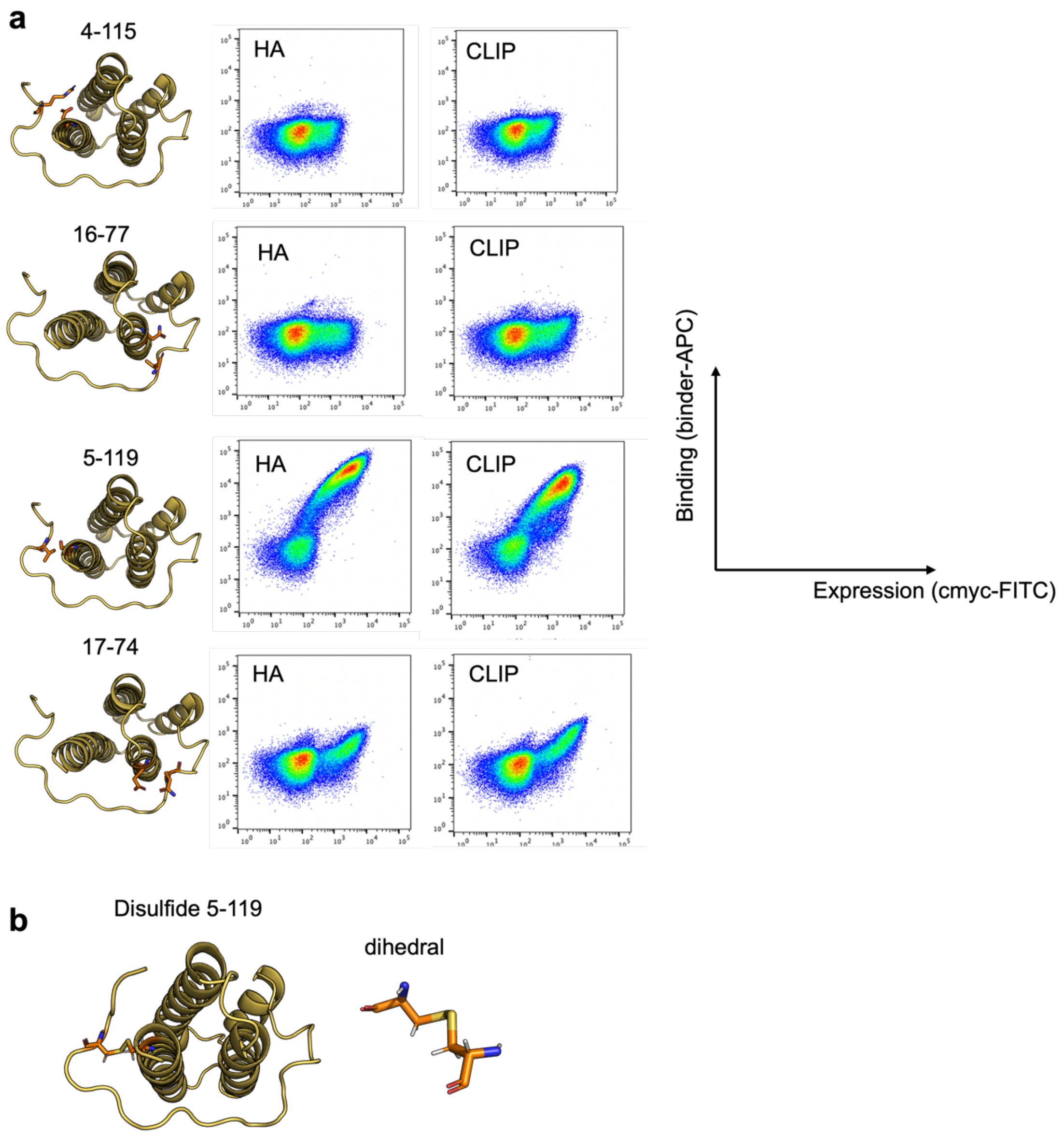


**Figure S3 Disulfide design**

A designed disulfide can attach the N-terminal loop onto the helical bundle, which restricts the loop flexibility and improves stability. **a.** All disulfide bonds given by RosettaRemodel that satisfy match_rt_limit < 1.1 were tested experimentally. A proper disulfide design should maintain the loop position and MHC tetramer binding signal. Among all 4 disulfide designs, the 5-119 disulfide gave the best binding signal. Left: Suggested disulfide positions. Right: binders displayed on the yeast surface and stained with pMHC tetramers. **B.** Left: model of 5-119 disulfide design. Right: dihedral view of disulfide and cysteine rotamers. Model gives a near 90-degree dihedral angle, indicating an energy-favored disulfide design.

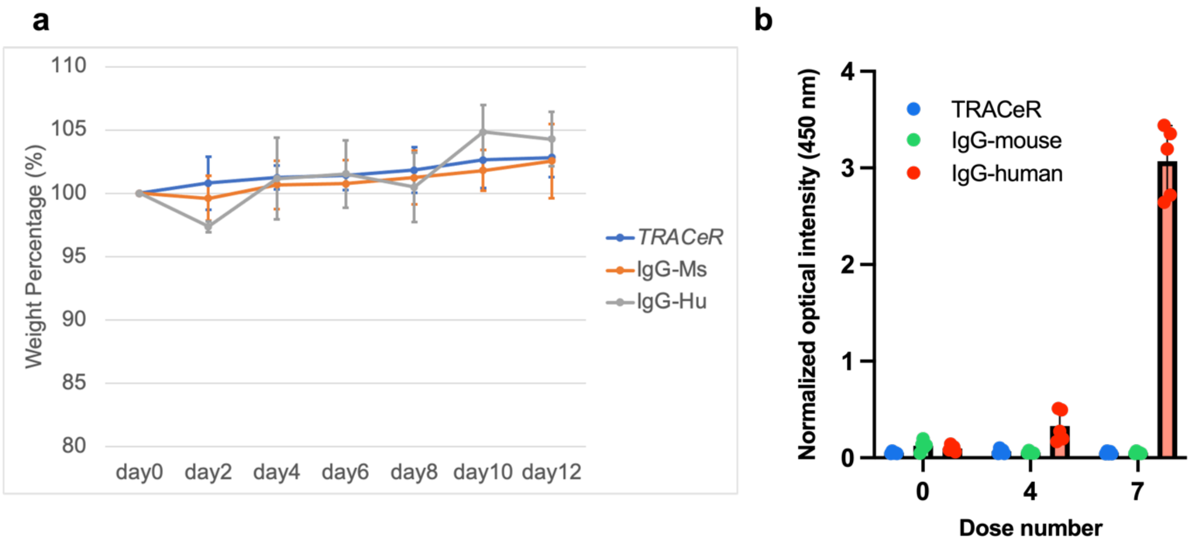


**Figure S4 $TRACeR^{CLIP}_{MHC-II,DR1}$ immunogenicity test (n=5)**

**a.** Mice weight change over the two-week injection period. **B.** Immune response in BALB/c mice (n=5) for mice that received 7 doses (0.2 mg/kg per dose) over two weeks. Serum was collected on day 0, day 7 and day 14. IgG responses before injection (dose 0), after 4 doses injection (dose 4), and after 7doses injection (dose 7) were measured by ELISA (1:500 serum).

HLA-DR1/ HA (PKYVKQNTLKLAT)

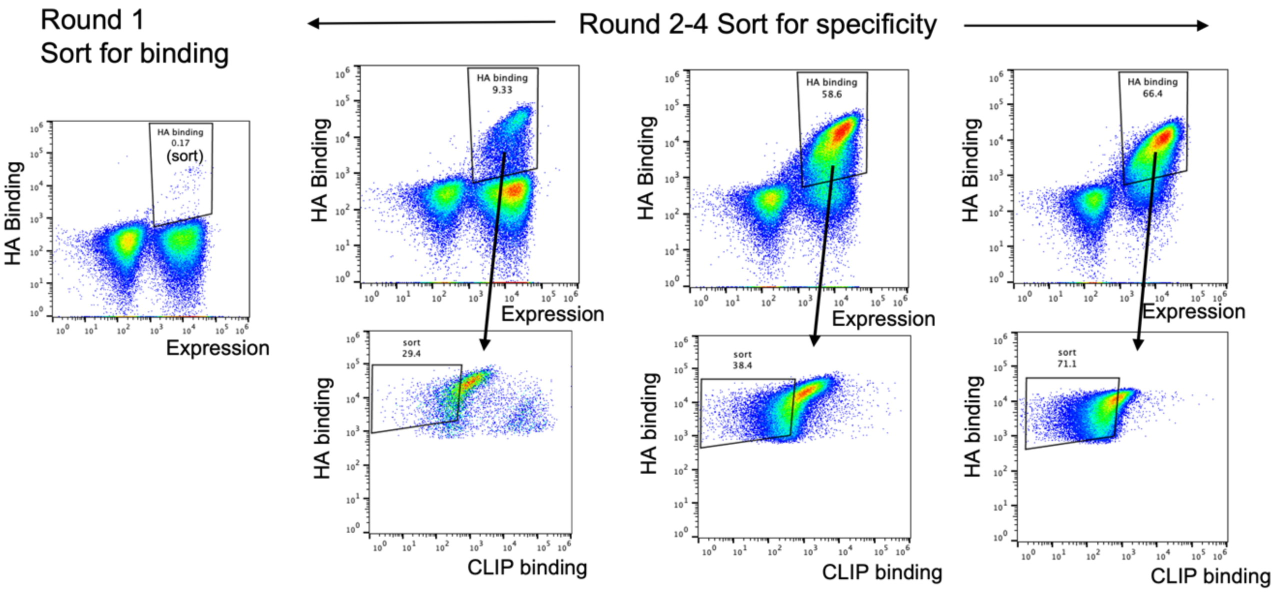


HLA-DR1/ NY-ESO-1(LLEFYLAMPFATPME)

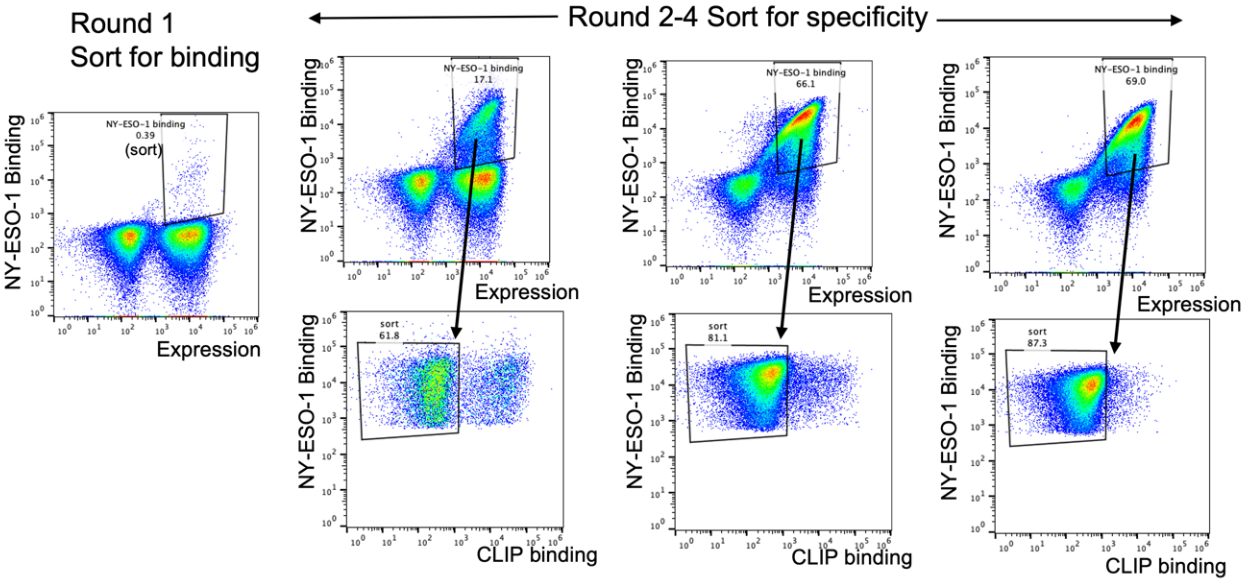

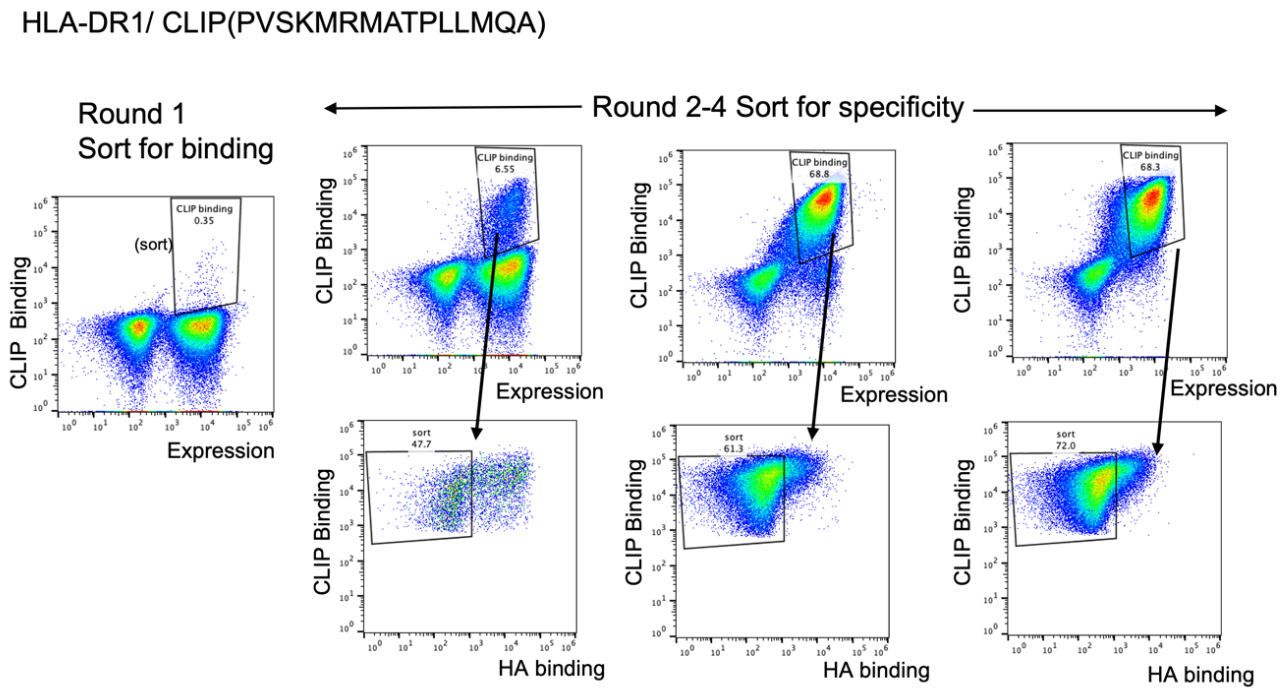


**Figure S5 Library selection for each target**

Library sorting and enrichment FACS plots. Sorting concentration is 50 nM tetramer for round1 and 50 nM tetramer for both on and off target for round2, and 25 nM tetramer for on target and 100 nM tetramer for off target starting from round3. Tetramers were provided by NIH tetramer core facility. In flow plot axis, expression level was profiled with anti-cmyc-FITC. HA binding was monitored with DR1/HA-APC. NY-ESO-1 binding was monitored with DR1/NY-ESO-1-APC. CLIP binding was monitored with DR1/CLIP-PE.

$TRACeR_{DR1}^{HA}$
(before optimization)

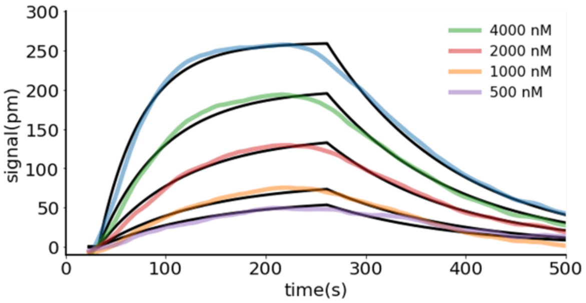


| | |
|---|---|
| $k_a$ (1/(M*s)) | 8.82e3 |
| $k_d$ (1/s) | 5.58e-3 |
| $K_D$ (M) | 6.32e-7 |

$TRACeR_{DR1}^{NY\text{-}ESO\text{-}1}$
(before optimization)

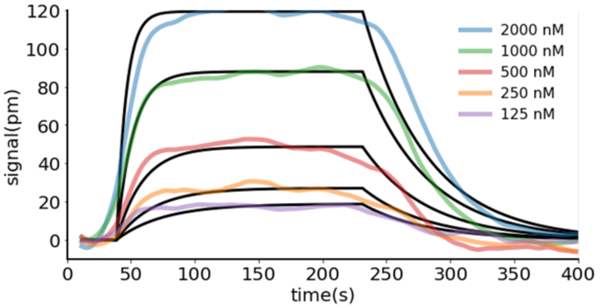


| | |
|---|---|
| $k_a$ (1/(M*s)) | 1.27e5 |
| $k_d$ (1/s) | 2.04e-2 |
| $K_D$ (M) | 1.60e-7 |

$TRACeR_{DR1}^{CLIP}$
(before optimization)

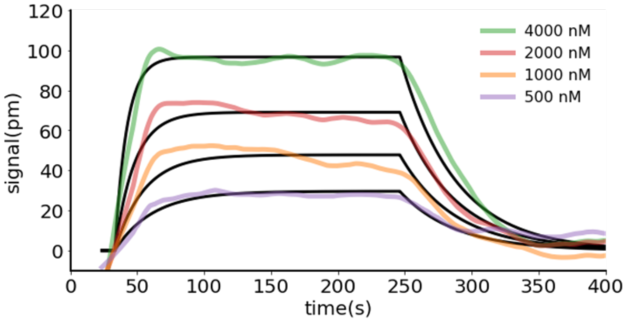


| | |
|---|---|
| $k_a$ (1/(M*s)) | 4.36e4 |
| $k_d$ (1/s) | 2.44e-2 |
| $K_D$ (M) | 5.60e-7 |

**Figure S6 *TRACeR-II* binding affinity with cognate antigens before affinity maturation by error-prone library**
Left: Binding kinetics measured with SPR. Right: Fitting parameters.

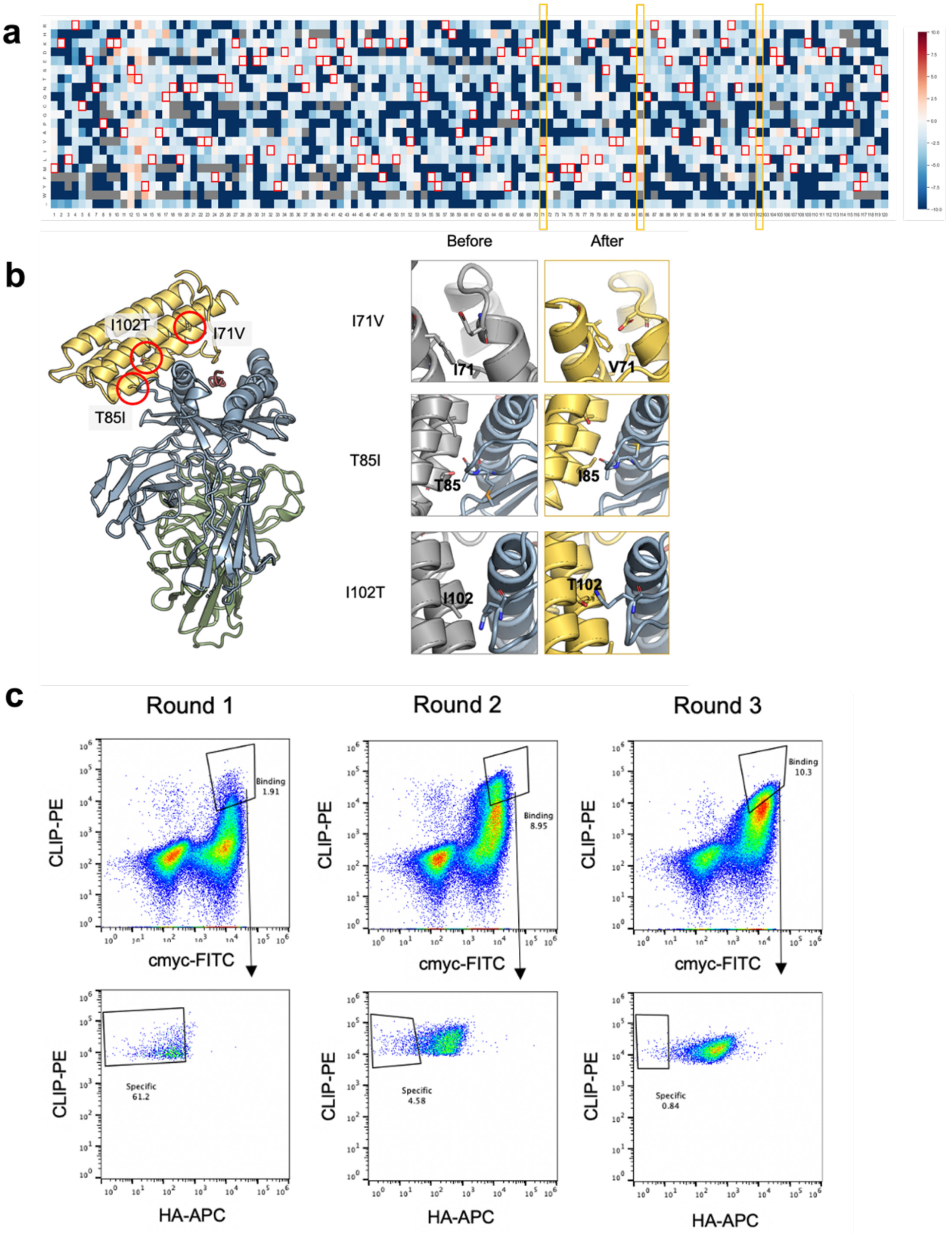


**Figure S7 Analysis of error-prone library for $TRACeR^{CLIP}_{MHC-II,DR1}$.**

**a.** Enrichment of each amino acid relative to the wildtype amino acid at each position. **b.** Structure comparison of native MAM and affinity matured scaffold on beneficial mutations. **c.** Selection process for binders with increased affinity and no off-target binding with yeast surface display.

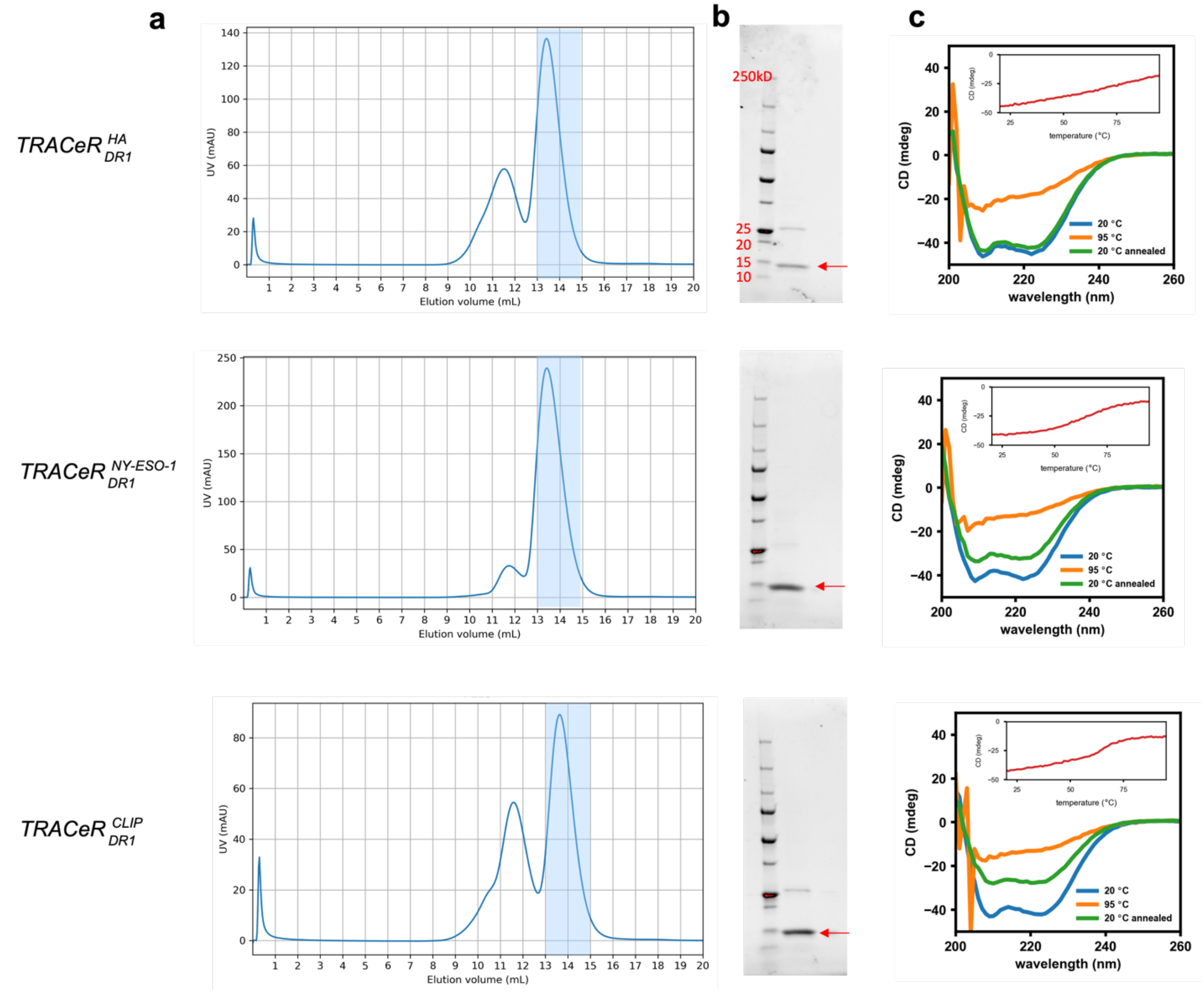


**Figure S8 Purification and biophysical characterization for $\boldsymbol{TRACeR^{HA}_{MHC-II,DR1}}$, $\boldsymbol{TRACeR^{NY-ESO-1}_{MHC-II,DR1}}$, $\boldsymbol{TRACeR^{CLIP}_{MHC-II,DR1}}$**

**a.** *TRACeR* protein purification from expression in *E.coli* BL21(DE3). All *TRACeR* proteins were highly soluble. The monomeric component was purified by SEC. Purification was performed on Superdex$^{TM}$ 75 column. Elution fractions in volume 13-14 mL were collected from SEC for downstream biophysical characterizations. **B.** Non-reducing SDS-PAGE for collected fractions. **C.** CD wavelength scanning and melting curve for purified *TRACeR*.

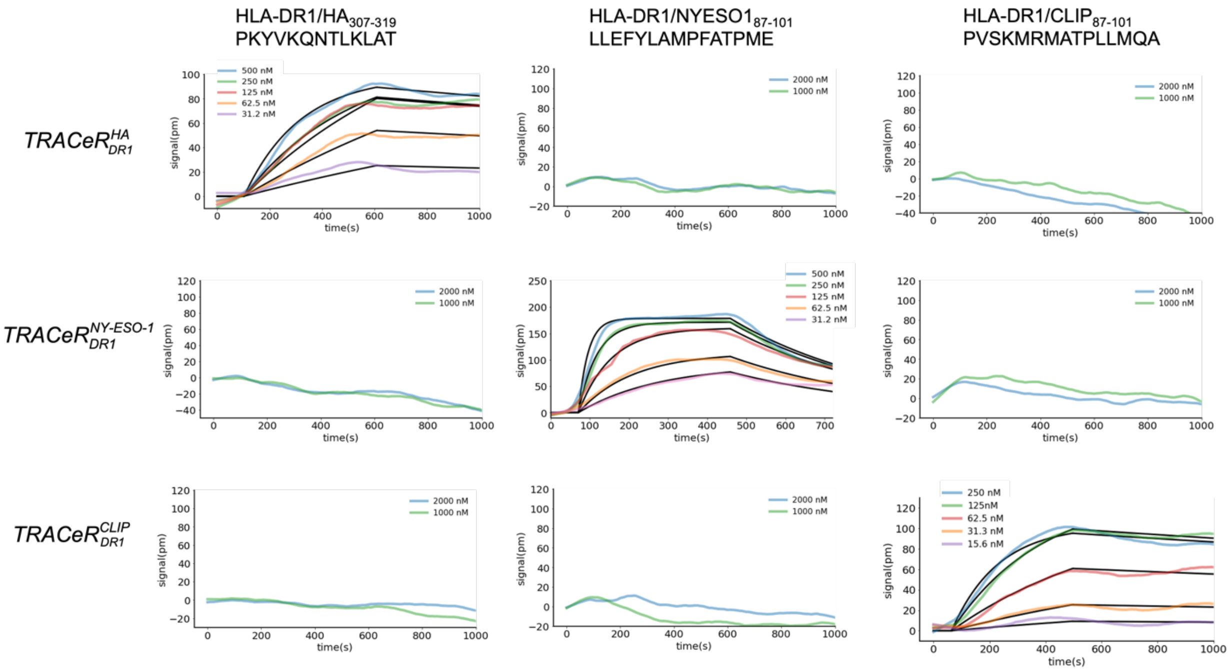


**Figure S9** ***TRACeR*** **binding specificity pattern as soluble proteins**
SPR measurement of *TRACeR* binding with on- and off-targets.

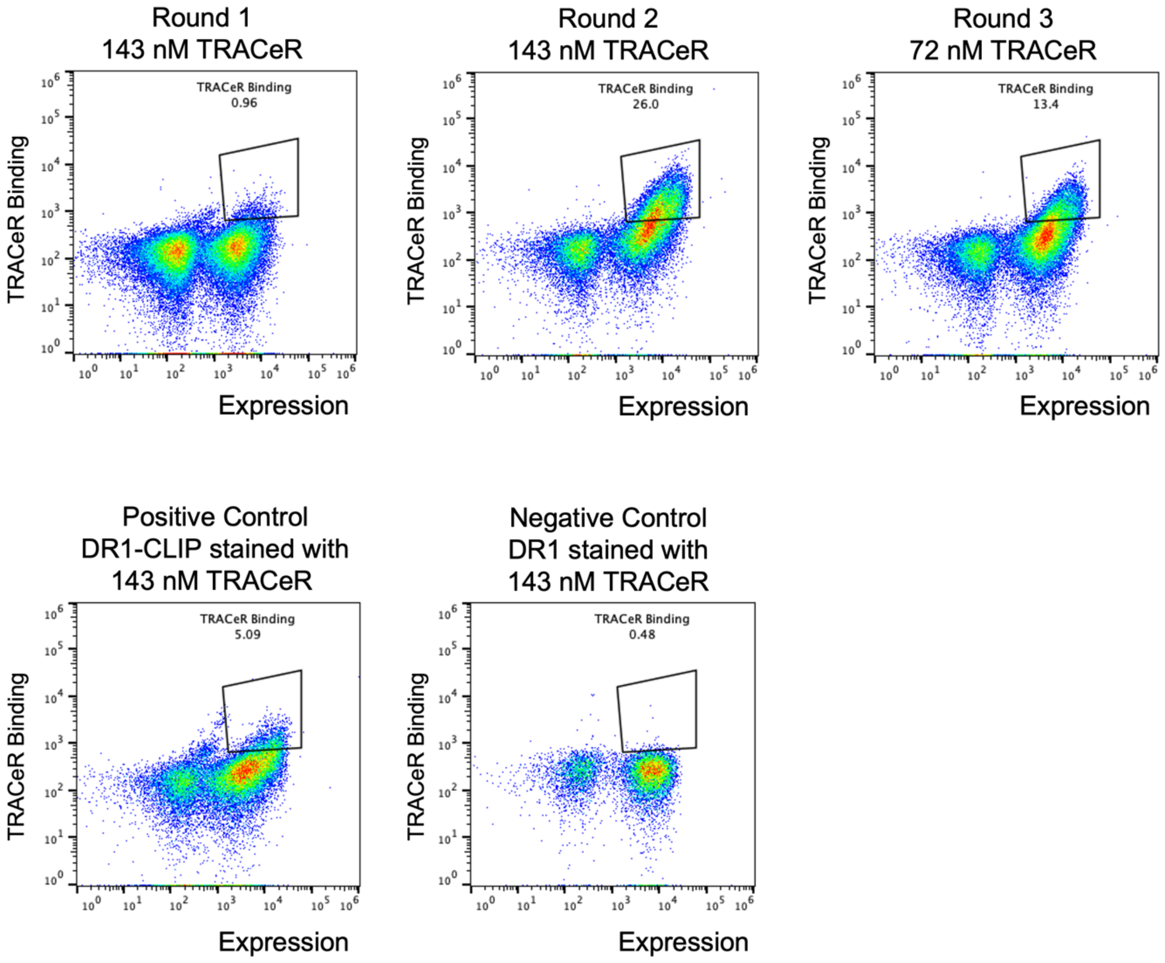


**Figure S10 Library sorting for determining $\boldsymbol{TRACeR^{CLIP}_{MHC-II,DR1}}$ specificity.**

Top: Selection of $TRACeR^{CLIP}_{MHC-II,DR1}$ binding peptide presented by HLA-DR1 from the SSM peptide library. *TRACeR* was biotinylated and co-incubated with streptavidin-PE for staining.

Bottom: *TRACeR* staining of yeast surface displayed HLA-DR1/CLIP (left) and HLA-DR1 without covalently linked peptide (right).

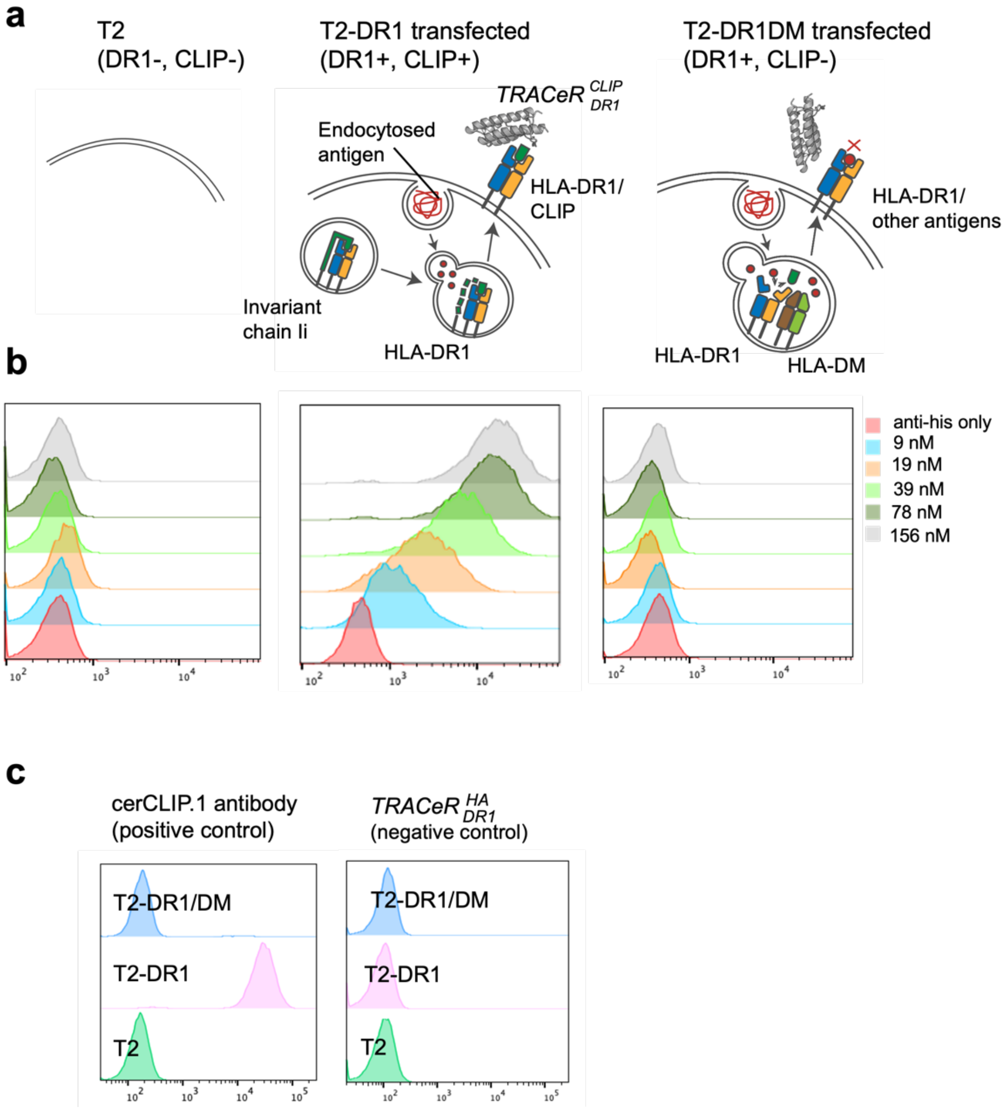


**Figure S11 *TRACeR* binds to cellularly processed antigens**

**a.** Schematic of T2 transfectants. **b.** Histogram of $TRACeR^{CLIP}_{MHC-II,DR1}$ staining T2 cell transfectants. **c.** Left: positive control with cerCLIP.1 antibody recognizing HLA-DR1/CLIP from a linear section outside of MHC-II binding groove (LPKPPKPVSKMRMATPLLMQALP). Right: negative control with $TRACeR^{HA}_{MHC-II,DR1}$.

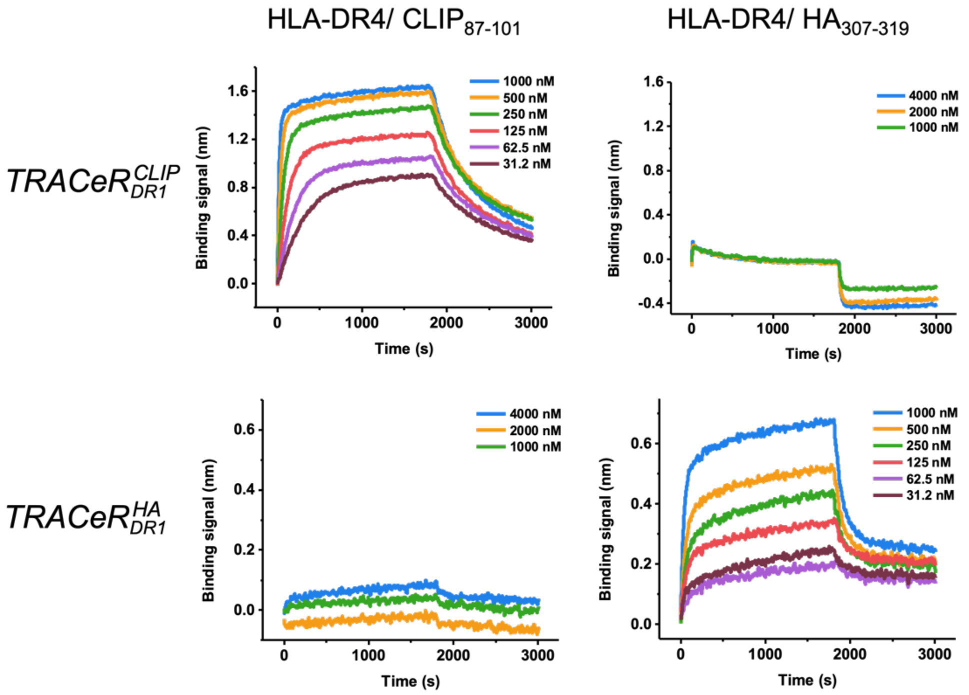


| | $K_D$ (M) | $K_D$ error | $k_{on}$ | $k_{on}$ error | $k_{dis}$ | $k_{dis}$ error |
|---|---|---|---|---|---|---|
| HLA-DR4/CLIP | 9.85e-8 | 2.31e-9 | 1.25e4 | 2.81e2 | 1.24e-3 | 8.63e-6 |
| HLA-DR4/HA | 4.09e-7 | 1.34e-8 | 6.73e3 | 1.87e2 | 2.76e-3 | 4.74e-5 |

**Figure S12 Binding and specificity pattern for DR1 *TRACeRs* on DR4-presented peptides**
Top: BLI measurement of $TRACeR^{CLIP}_{MHC-II,DR1}$ binding to HLA-DR4/CLIP and HLA-DR4/HA.
Middle: BLI measurement of $TRACeR^{HA}_{MHC-II,DR1}$ binding to HLA-DR4/CLIP and HLA-DR4/HA.
Bottom: Fitting parameters of binding kinetics. Data were analyzed and processed using ForteBio Data Analysis software 7.1.0.100.

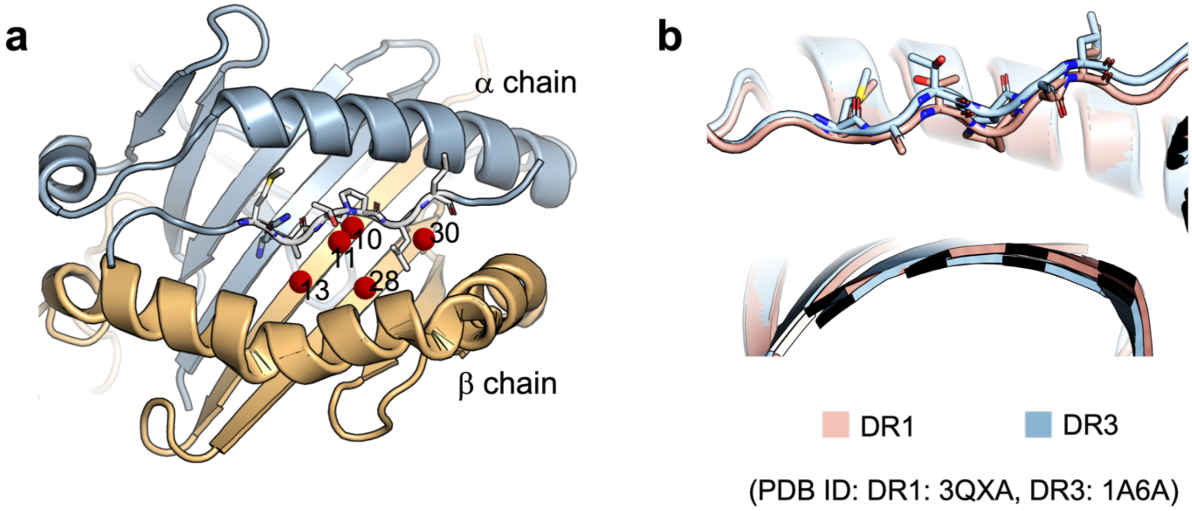


c

| Position \ Allele | DR1 | DR4 | DR3 | DR15 |
|---|---|---|---|---|
| 10 | W | E | Y | Q |
| 11 | L | V | S | D |
| 13 | F | H | S | Y |
| 28 | E | D | D | H |
| 30 | C | Y | Y | D |

(PDB ID: DR1: 3QXA, DR4: 1J8H, DR3: 1A6A, DR15:1H15)

**Figure S13. Polymorphic β chain on DR alleles affects presented peptide conformation**
The polymorphism of HLA-DR alleles results in different side chain identities in the peptide binding pocket, which further leads to different peptide conformations. **a.** The peptide binding pocket is polymorphic among different DR alleles, resulting in different peptide conformations associated with the same sequence. Red spheres: polymorphic positions on β chain. **B.** The CLIP peptide conformation difference between HLA-DR1 and HLA-DR3 in crystal structures. T9 from CLIP peptide forms a critical hydrogen bond interaction with the ARE region of *TRACeR*. However, A significant position shift on the T9 from CLIP peptide is observed between DR1 and DR3 due to the sequence diversity from the supporting residues in peptide binding groove. **C.** Amino acid identity at polymorphic positions in the peptide binding pocket.

HLA-DR3/ HER2(SANIQEFAGCKKIFG )

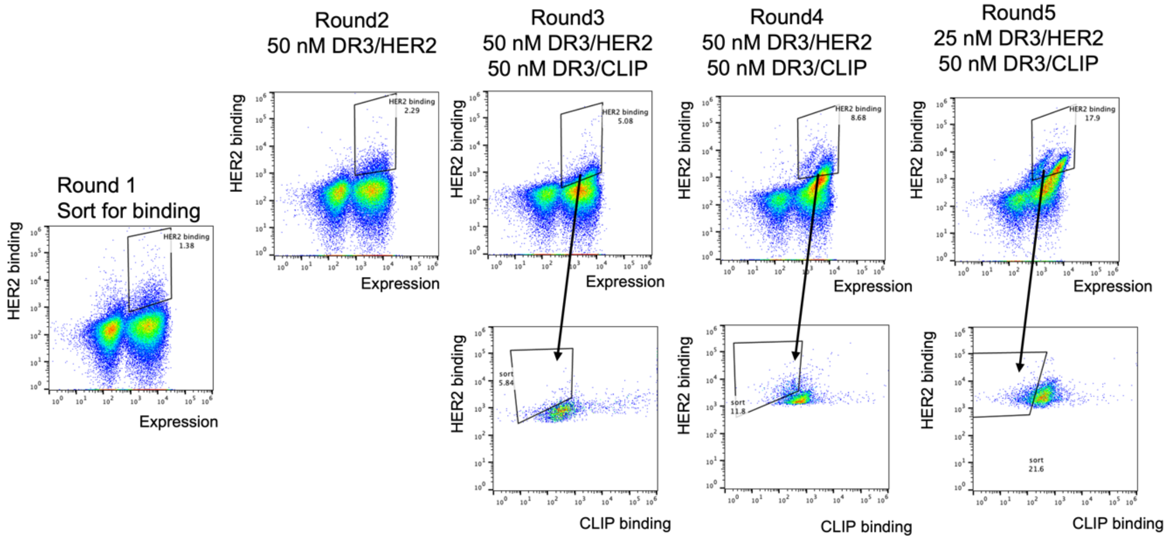


HLA-DR15/αSynuclein (KTKEGVLYVGSKTKE)

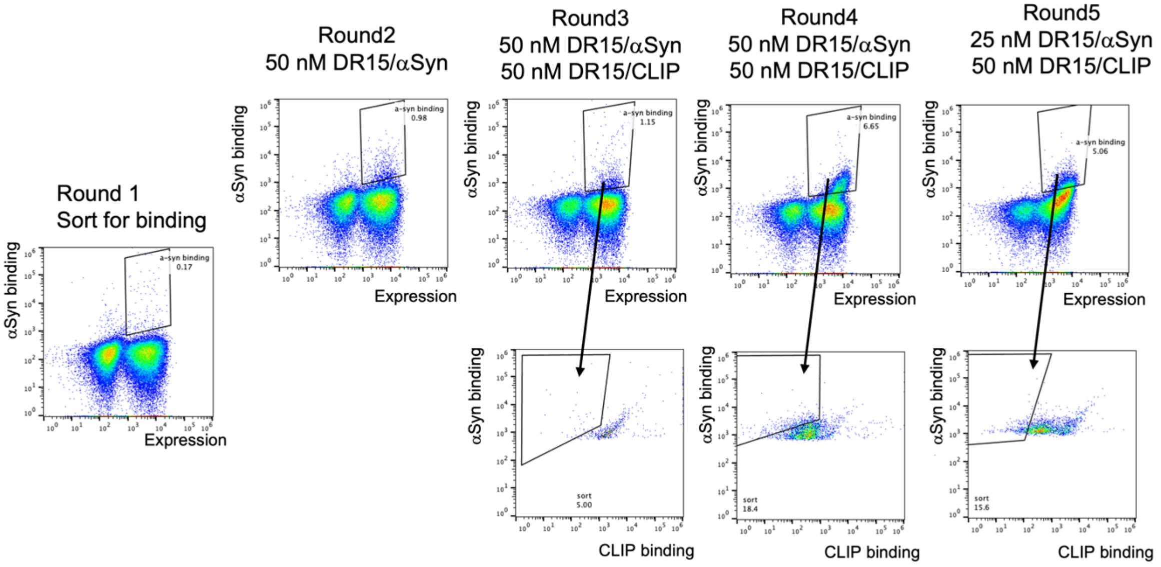

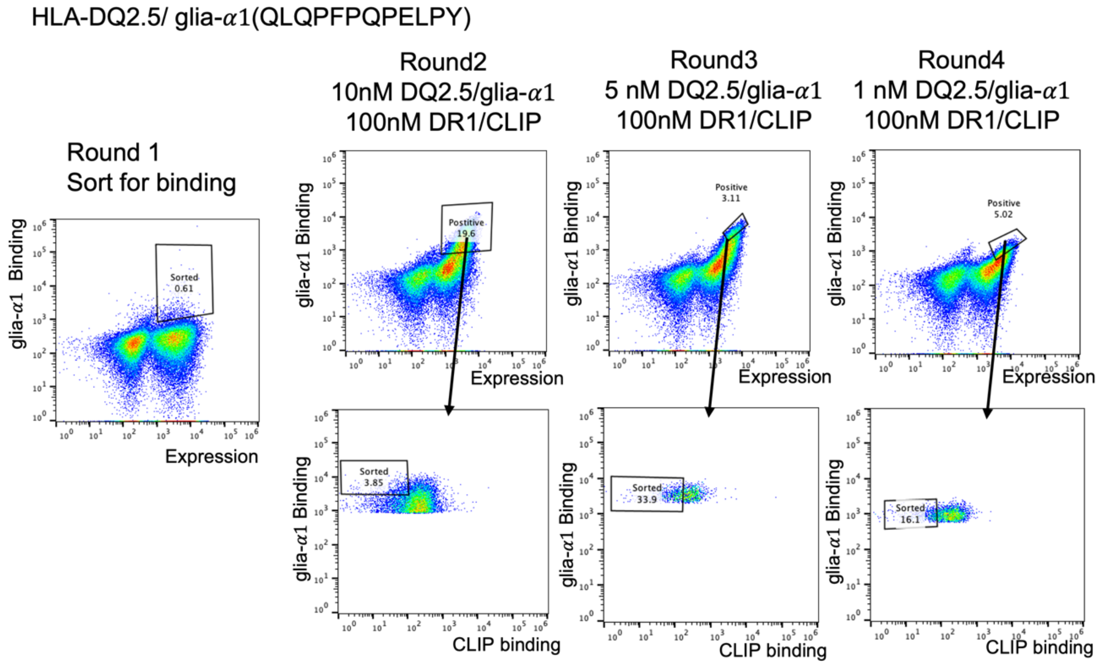


**Figure S14 Library sorting for $TRACeR^{HER2}_{MHC-II,DR3}$ , $TRACeR^{\alpha Syn}_{MHC-II,DR15}$ , $TRACeR^{glia-\alpha 1a}_{MHC-II,DQ2.5}$ development**

Library sorting and enrichment FACS plots. All concentrations are corresponding MHC-II tetramer concentrations.

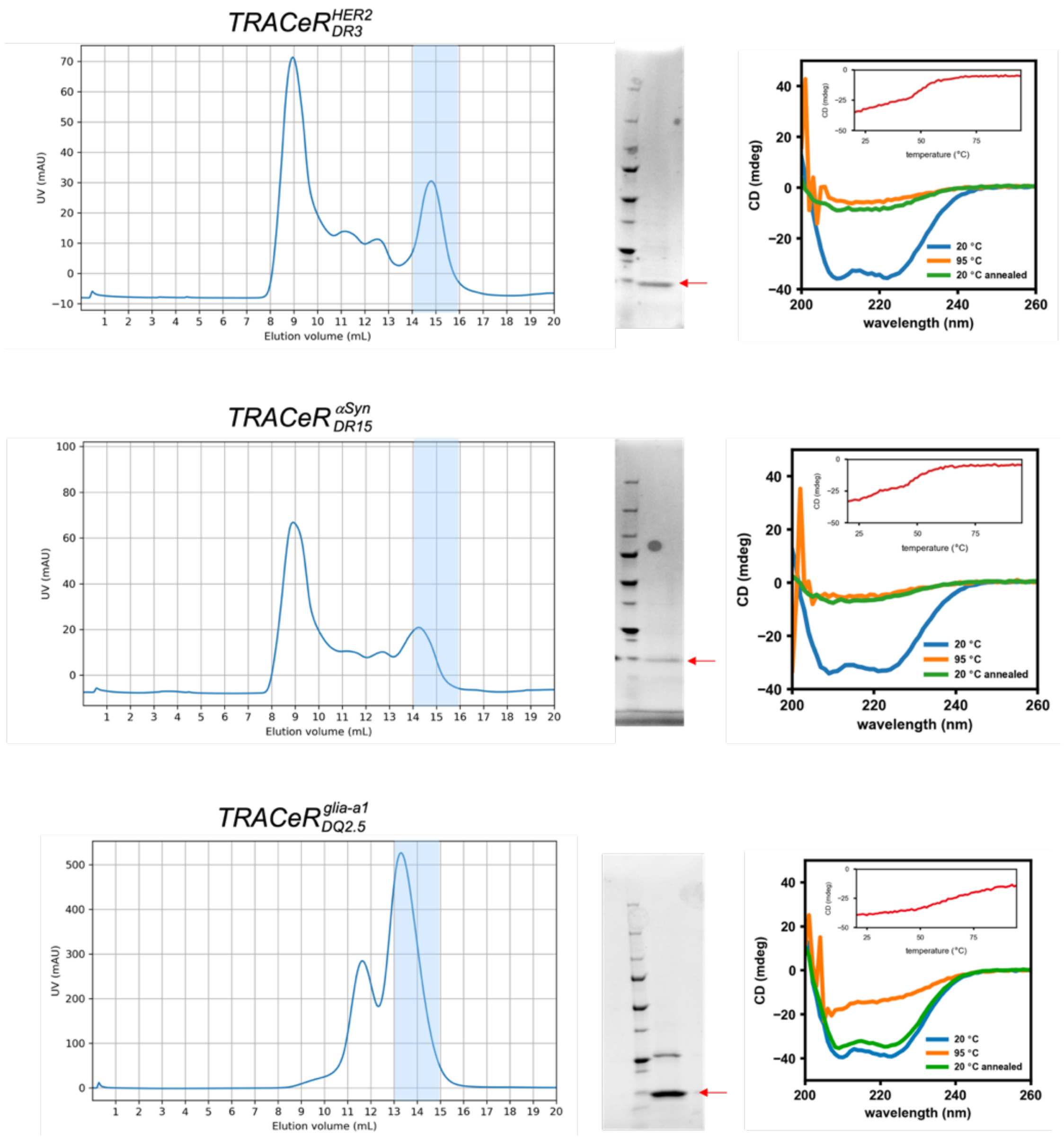


**Figure S15 Purification and biophysical characterization of $\boldsymbol{TRACeR^{HER2}_{MHC-II,DR3}}$, $\boldsymbol{TRACeR^{aSyn}_{MHC-II,DR15}}$, $\boldsymbol{TRACeR^{glia-a1}_{MHC-II,DQ2.5}}$.**

Left: The monomeric component was purified by SEC. $TRACeR^{HER2}_{MHC-II,DR3}$ and $TRACeR^{aSyn}_{MHC-II,DR15}$ were purified on Superdex$^{TM}$ 75 increase column. $TRACeR^{glia-a1}_{MHC-II,DQ2.5}$ was purified on Superdex$^{TM}$ 75 column. Elution fractions colored in blue were collected from SEC for downstream characterization.

Middle: Non-reducing SDS-PAGE for the collected fractions.

Right: CD wavelength scanning and melting curve for purified *TRACeR*.

Error-prone mutations are not incorporated.

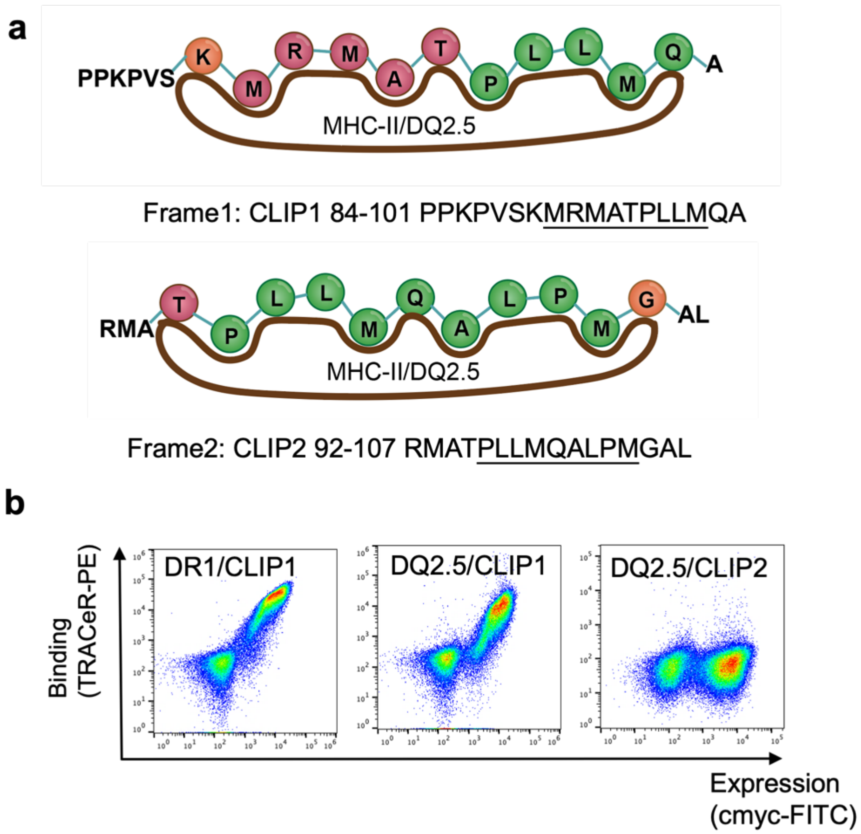


**Figure S16 Alternative presentation frames for CLIP peptide on HLA-DQ2.5 allele and *TRACeR* binding**

**a.** Schematic of alternative antigen presentation frames. **B.** *TRACeR* binding to HLA-DR1/CLIP1, HLA-DQ2.5/CLIP1, HLA-DQ2.5/CLIP2. *TRACeR* was expressed on the yeast surface and stained with 50 nM pMHC tetramer.

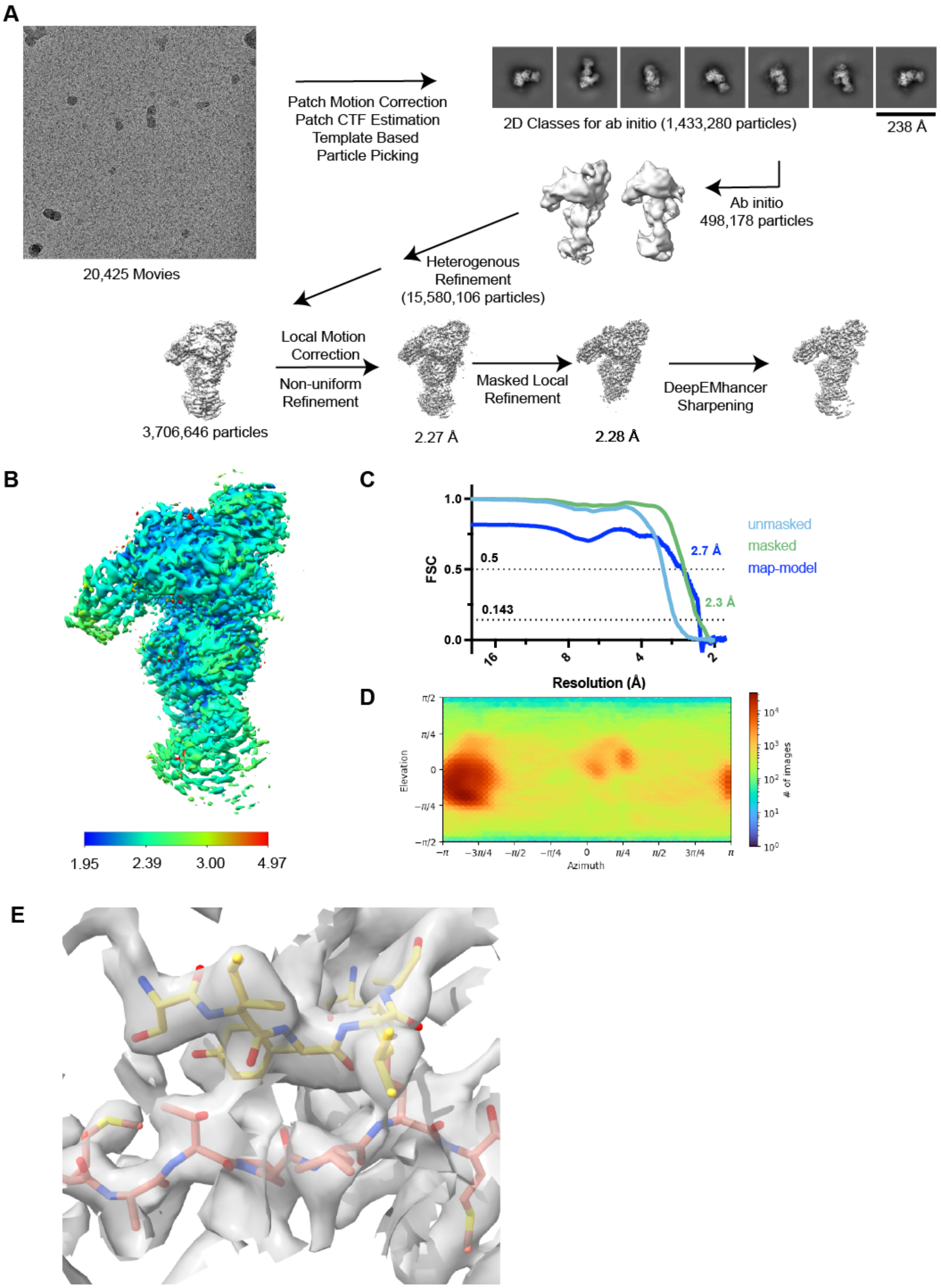


**Figure S17 Data processing for $TRACeR^{CLIP}_{MHC-II,DR1}$- HLA-DR1/CLIP-c44H10 Fab ternary complex**

**a.** The workflow for cryoEM data processing. **B.** Local-refined map colored by resolution **c.** Gold Standard Fourier Shell Coefficient resolution of the local-refined map. **D.** Distribution of different particle orientations **e.** ARE-peptide interface shown with the DeepEMHancer-sharpened map; ARE is shown in yellow and the peptide in salmon.

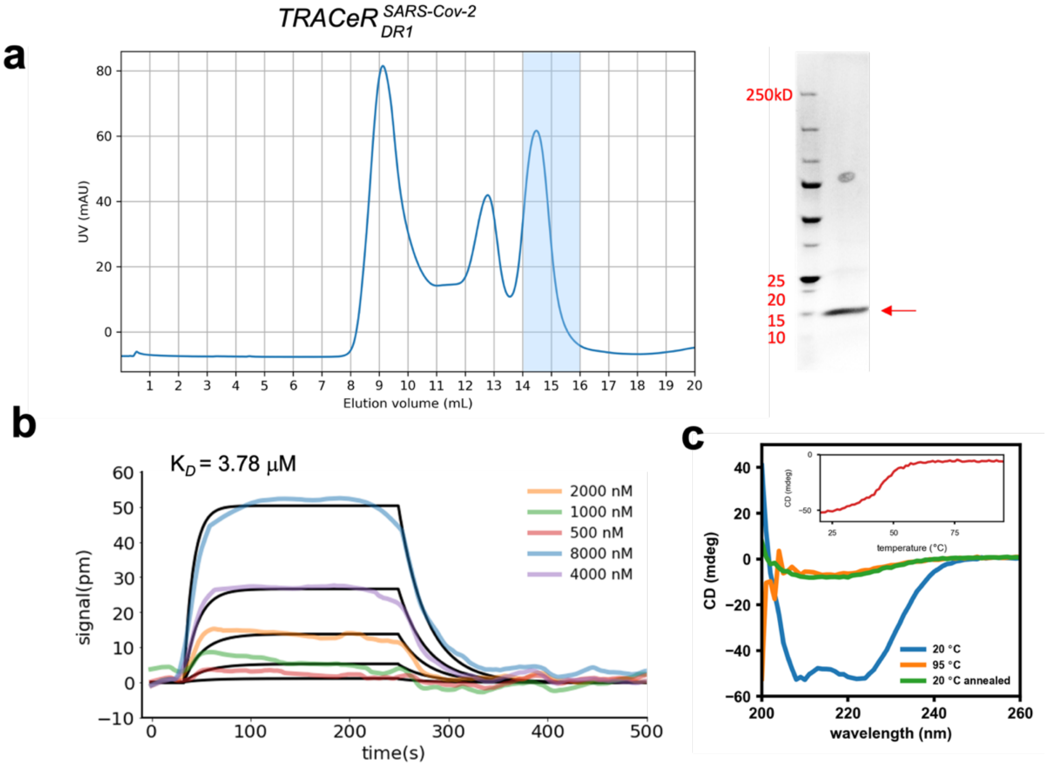


**Figure S18 Purification and binding affinity measurement of designed $TRACeR^{SARS-CoV-2}_{MHC-II,DR1}$**
**a.** SEC purification curve and SDS-PAGE (non-reducing) of purified *TRACeR.* Purification was performed on Superdex$^{TM}$ 75 increase column. Fractions 14 and 15 were collected for further characterization. **b.** Binding kinetics determined by SPR measurement. **c.** CD curve of the purified monomer fraction.

**Supplementary table 1 Summary of all relevant amino acid sequences**

Sequences will be released upon publication

**Supplementary table 2. Binding kinetics summary of *TRACeRs***

| *TRACeR* | $K_D$ (M) | $K_D$ error | $k_{on}$ (1/(M*s)) | $k_{on}$ error | $k_{dis}$ (1/s) | $k_{dis}$ error |
|---|---|---|---|---|---|---|
| HLA-DR1/CLIP | 8.02e-9 | 1.70e-10 | 2.33e4 | 1.64e1 | 1.87e-4 | 3.86e-6 |
| HLA-DR1/HA | 4.76e-8 | 6.10e-10 | 4.50e3 | 1.24e1 | 2.14e-4 | 2.16e-6 |
| HLA-DR1/NY-ESO-1 | 3.29e-8 | 4.02e-10 | 4.38e4 | 8.48e1 | 1.44e-3 | 1.48e-5 |
| HLA-DR3/HER2 | 2.32e-6 | 2.95e-8 | 7.25e3 | 9.17e1 | 1.68e-2 | 7.95e-7 |
| HLA-DR15/αSyn | 1.69e-5 | 2.21e-6 | 7.93e2 | 1.02e2 | 1.34e-2 | 1.67e-6 |
| HLA-DQ2.5/glia-α1 | 4.61e-7 | 8.71e-8 | 4.74e4 | 8.48e3 | 2.19e-2 | 1.33e-3 |
| HLA-DR1/SARS-CoV-2 | 3.31e-7 | 1.50e-10 | 5.08e4 | 2.23e1 | 1.68e-2 | 2.29e-7 |

**Supplementary table 3. Cryo-EM data processing**

| **Data collection and processing** | |
|---|---|
| Nominal magnification | 165,000 |
| Acceleration voltage (kV) | 300 |
| Electron exposure ($e^-/Å^2$) | 60 |
| Defocus range (μm) | -0.8 – -1.8 |
| Pixel size (Å) | 0.743 |
| Symmetry imposed | C1 |
| Final particle images | 3,706,646 |
| Map resolution FSC threshold | 0.143 |
| Map resolution (Å) | 2.34 |
| **Refinement** | |
| Initial model used (PDB) | 6ROE, 8EUQ, 2ICW |
| Model resolution FSC threshold | 0.5 |
| Model resolution (Å) | 2.69 |
| Model composition | |
| Non-hydrogen atoms | 5835 |
| Protein residues | 718 |
| Ligands | 3 |
| B-factors ($Å^2$) | |
| Protein | 71.48 |
| Ligand | 100.08 |
| R.M.S. deviations | |
| Bond lengths (Å) | 0.002 |
| Bond angles (°) | 0.461 |
| **Validation** | |
| Molprobity score | 1.61 |
| Clashscore | 6.78 |
| Rotamer outliers (%) | 1.71 |
| Ramachandran plot | |
| Favored (%) | 97.73 |
| Allowed (%) | 2.27 |
| Outliers (%) | 0 |

**Supplementary Appendix 1. Design workflow for $TRACeR^{SARS-CoV-2}_{MHC-II,DR1}$**

1. Starting structure preparation
The crystal structure of superantigen MAM binding with HLA-DR1 presenting HA peptide was downloaded from PDB (PDB ID: 2ICW). Chain A (MHC α chain), B (MHC β chain), C (peptide), G (MAM) are extracted from the structure and cleaned to remove modalities not recognized by Rosetta (clean_pdb.py in Rosetta suite).
The structure containing chain A, B, C, G is then relaxed under constrain for further Rosetta design. The PDB file is further modified to have chain order ABGC for simplicity in following steps.
rosetta.binary.m1.release-340/main/source/bin/relax.static.macosclangrelease -relax:constrain_relax_to_start_coords -relax:coord_constrain_sidechains -relax:ramp_constraints false -s 2icw_ABCG.pdb -ex1 -ex2 -use_input_sc -flip_HNQ -no_optH false

1. Peptide threading

Binding frame of the SARS-CoV-2 antigen GAALQIPFAMQMAYRF is predicted by NetMHCIIpan-3.2 online server (https://services.healthtech.dtu.dk/services/NetMHCIIpan-3.2/) to have the core binding frame of FAMQMAYRF. The SARS-CoV-2 antigen is threaded onto the HA antigen using Rosetta fixbb application, the C terminal end of the HA antigen is truncated to fit the new antigen length.

2icw_ABCG_0001_ABGC_to_SARS2.resfile

```
NATRO
START

577 C PIKAA I
578 C PIKAA P
579 C PIKAA F
580 C PIKAA A
581 C PIKAA M
582 C PIKAA Q
583 C PIKAA M
584 C PIKAA A
585 C PIKAA Y
586 C PIKAA R
587 C PIKAA F
```

Command:

rosetta.binary.m1.release-340/main/source/bin/fixbb.static.macosclangrelease -s 2icw_ABCG_0001_ABGC_short.pdb -resfile 2icw_ABCG_0001_ABGC_to_SARS2.resfile -ex1 -ex2 -nstruct 1 -out:prefix SARS2

2. Binder Design

To facilitate the interaction between MAM and the peptide, the behavior of the ProteinProteinInterfaceUpweighter Mover was modified by removing the "sec=L" (line 105 and line 106) in /main/source/src/protocols/toolbox/IGEdgeReweighters.cc and recompile the rosetta_script application. The modification allows only upweighting intereaction between chain C and G.

The binder was designed by using the following Rosetta script. The script first forms the critical disulfide bond, then design the MAM binding loop in 2 FastDesign and FastRelax cycles. 1000 designs are generated.

rosetta.source.release-340/main/source/bin/rosetta_scripts.linuxgccrelease -s SARS22icw_ABCG_0001_ABGC_short_0001.pdb -use_input_sc -nstruct 10 -ex1 -ex2 -parser:protocol fastdesign_2upweight_interface_DRDR_dsf.xml -corrections::beta_nov16 -precompute_ig -out:prefix R04_w2_$SLURM_ARRAY_TASK_ID_

3. Scoring

Structures are scored by alphafold2. The average pLDDT of the 5 loop residues of each design is used to rank the structures. Top 10 ranked designs (excluding repeats) are used for experimental validation.

**Supplementary Appendix 2. Script for designing *TRACeR* ARE region**

```
<ROSETTASCRIPTS>

# Adapted from Brian Coventry and Longxing Cao 2020


  <SCOREFXNS>
    <ScoreFunction name="sfxn" weights="beta_nov16" />
    <ScoreFunction name="sfxn_relax" weights="beta_nov16" >
      <Reweight scoretype="arg_cation_pi" weight="3" />
      <Reweight scoretype="approximate_buried_unsat_penalty" weight="5" />
      <Set approximate_buried_unsat_penalty_burial_atomic_depth="3.5" />
      <Set approximate_buried_unsat_penalty_hbond_energy_threshold="-0.5" />
    </ScoreFunction>
    <ScoreFunction name="sfxn_design" weights="beta_nov16" >
      <Reweight scoretype="res_type_constraint" weight="1.5" />
      <Reweight scoretype="aa_composition" weight="1.0" />
      <Reweight scoretype="arg_cation_pi" weight="3" />
      <Reweight scoretype="approximate_buried_unsat_penalty" weight="5" />
      <Set approximate_buried_unsat_penalty_burial_atomic_depth="3.5" />
      <Set approximate_buried_unsat_penalty_hbond_energy_threshold="-0.5" />
      <Set approximate_buried_unsat_penalty_hbond_bonus_cross_chain="-1" />
    </ScoreFunction>
```

```
  </SCOREFXNS>

  <RESIDUE_SELECTORS>
      <Chain name="chainA" chains="A"/> # DR1 alpha chain
      <Chain name="chainB" chains="B"/> # DR1 beta chain
      <Chain name="chainC" chains="C"/> # peptide
      <Chain name="chainG" chains="G"/> # MAM
      <Or name="DR1" selectors="chainA, chainB" />
      <Or name="DR1_side" selectors="chainA, chainB, chainC" />
      <Index name="loop_5" resnums="375,376,377,378,379" /> # ARE loop
      <Neighborhood name="neighbors_loop_ex" selector="loop_5" distance="12.0"
include_focus_in_subset="False" />
      <Neighborhood name="neighbors_loop_in" selector="loop_5" distance="12.0"
include_focus_in_subset="True" />
      <Neighborhood name="neighbors_antigen_in" selector="chainC" distance="10.0"
include_focus_in_subset="True" />
      <And name="neighbors_antigen_MHC" selectors="DR1, neighbors_antigen_in" />
      <And name="neighbors_antigen_MAM" selectors="chainG, neighbors_antigen_in" />
      <Or name="everything" selectors="chainA, chainB, chainC, chainG" />
      <Not name="Not_neighbor" selector="neighbors_loop_in" />
      <And name="background" selectors="everything, Not_neighbor" />
      <Index name="disulfide" resnums="482,368" />
      <Neighborhood name="around_disulfide" selector="disulfide" distance="12.0"
include_focus_in_subset="True" />
      <Not name="Not_around_disulfide" selector="around_disulfide" />
      <And name="background_disulfide" selectors="everything, Not_around_disulfide" />
  </RESIDUE_SELECTORS>

  <TASKOPERATIONS>

      <PruneBuriedUnsats name="prune_buried_unsats" allow_even_trades="false"
atomic_depth_cutoff="3.5" minimum_hbond_energy="-0.5" />

      <ProteinProteinInterfaceUpweighter name="upweight_interface" interface_weight="2"
skip_loop_in_chain="A,B" /> # Very important, only upweight interaction with peptide
      <ProteinInterfaceDesign name="pack_long" design_chain1="0" design_chain2="0"
jump="1" interface_distance_cutoff="15"/>
      <IncludeCurrent name="current" />
      <LimitAromaChi2 name="limitchi2" chi2max="110" chi2min="70" include_trp="True" />
      <ExtraRotamersGeneric name="ex1_ex2" ex1="1" ex2aro="1" ex2="0" />

      <OperateOnResidueSubset name="not_around_disulfide_freeze"
selector="background_disulfide">
          <PreventRepackingRLT/>
      </OperateOnResidueSubset>
      <OperateOnResidueSubset name="repacking_disulfide" selector="around_disulfide">
```

```
            <RestrictToRepackingRLT/>
        </OperateOnResidueSubset>

        <OperateOnResidueSubset name="background_no_packing" selector="background">
            <PreventRepackingRLT/>
        </OperateOnResidueSubset>
        <OperateOnResidueSubset name="neighbors_repacking_only"
selector="neighbors_loop_ex">
            <RestrictToRepackingRLT/>
        </OperateOnResidueSubset>



    </TASKOPERATIONS>

    <FILTERS>
    </FILTERS>

    <SIMPLE_METRICS>
    </SIMPLE_METRICS>

    <MOVERS>

        <MutateResidue name="mutate_1st" target="482G" new_res="CYS"
preserve_atom_coords="false" mutate_self="false" update_polymer_bond_dependent="false" />
        <MutateResidue name="mutate_2nd" target="368G" new_res="CYS"
preserve_atom_coords="false" mutate_self="false" update_polymer_bond_dependent="false" />
        <ForceDisulfides name="form_disulfide" scorefxn="sfxn" disulfides="482G:368G"
remove_existing="false" repack="true" />
        <FastDesign name="FastDesign" scorefxn="sfxn_design" repeats="1"
task_operations="current,limitchi2,ex1_ex2,background_no_packing,neighbors_repacking_only,
upweight_interface" batch="false" ramp_down_constraints="false" cartesian="false"
bondangle="false" bondlength="false" min_type="dfpmin_armijo_nonmonotone"
relaxscript="/scratch/users/ljjchem/MHCII_loopdesign/old_beta_16_ref.rosettacon2018.txt" >
            <MoveMap name="MM" bb="false" chi="false" jump="false" >
                    <ResidueSelector selector="loop_5" chi="true" bb="true" /> # Redesign ARE allow
both backbone and sidechain movement
                    <ResidueSelector selector="neighbors_loop_ex" chi="true" bb="true" /> # Allow
neighbor residues to repack without design
                    <Jump number="3" setting="true" />
            </MoveMap>
        </FastDesign>

        <FastRelax name="FastRelax" scorefxn="sfxn_design" repeats="1" batch="false"
ramp_down_constraints="false" cartesian="false" bondangle="false" bondlength="false"
min_type="dfpmin_armijo_nonmonotone" task_operations="ex1_ex2,limitchi2" >
```

```
        <MoveMap name="MM" bb="false" chi="false" jump="false" >
            <ResidueSelector selector="neighbors_antigen_MHC" chi="true" bb="false" />
            <ResidueSelector selector="loop_5" chi="true" bb="true" /> #
            <ResidueSelector selector="neighbors_loop_ex" chi="true" bb="true" />
            <ResidueSelector selector="chainC" chi="true" bb="true" />
            <Jump number="3" setting="true" />
            <Jump number="2" setting="true" />
        </MoveMap>
      </FastRelax>

      <FastRelax name="relax_disulfide" scorefxn="sfxn_design" repeats="1" batch="false"
ramp_down_constraints="false" cartesian="false" bondangle="false" bondlength="false"
min_type="dfpmin_armijo_nonmonotone"
task_operations="ex1_ex2,limitchi2,not_around_disulfide_freeze,repacking_disulfide" >

        <MoveMap name="MM" bb="false" chi="false" jump="false" >
            <ResidueSelector selector="around_disulfide" chi="true" bb="true" />
        </MoveMap>
      </FastRelax>


    </MOVERS>
    <APPLY_TO_POSE>
    </APPLY_TO_POSE>
    <PROTOCOLS>
      # Form disulfide
      <Add mover="mutate_1st" />
      <Add mover="mutate_2nd" />
      <Add mover="form_disulfide" />
      <Add mover="relax_disulfide" />
      # perform sequence optimization
      <Add mover="FastDesign" />
      <Add mover="FastRelax" />
      <Add mover="FastDesign" />
      <Add mover="FastRelax" />

    </PROTOCOLS>
</ROSETTASCRIPTS>
```
